# Carbon source and surface type influence the early-stage biofilm formation by rhizosphere bacterium *Pseudomonas donghuensis* P482

**DOI:** 10.1101/2023.06.30.547207

**Authors:** Magdalena Rajewska, Tomasz Maciąg, Sylwia Jafra

## Abstract

The competence of bacteria to colonize different environmental niches is often determined by their ability to form biofilms. This depends on both cellular and extracellular factors, such as individual characteristics of a strain, type of colonized surface (abiotic or biotic) or availability and source of nutrients. *Pseudomonas donghuensis* P482 efficiently colonizes rhizosphere of various plant hosts, but a connection between plant tissue colonization and biofilm formation has not been verified for P482 up to date. Here we demonstrate that the ability of P482 to form biofilm on abiotic surfaces and the structural characteristics of the biofilm are connected to the type of carbon source available to the bacteria, with glycerol promoting formation of developed biofilm at early stages. Also, the type of substratum, polystyrene or glass, significantly influences the ability of P482 to attach to the surface, possibly due to hydrophobic effects. Moreover, mutants in genes associated with motility or chemotaxis, synthesis of polysaccharides, and encoding proteases or regulatory factors, affected in biofilm formation on glass were fully capable of colonizing root tissue of both tomato and maize hosts. This indicates that the ability to form biofilm on distinct abiotic surfaces does not simply correlate with the efficient colonization of rhizosphere and formation of biofilm on plant tissue by P482.

## Introduction

Biofilm is the most frequently encountered form of bacterial growth in the majority of ecosystems, estimating even up to 80% of cells living in community-like gatherings (Flemming and Wuertz, 2019). Bacterial cells building the biofilms may interact with both biotic and abiotic surfaces, creating complex, multicellular structures which guarantee survival in specific environmental niches through protection from nutrient depletion, activity of antibiotics and bactericides, harmful influence of the environment, predation or simply physical damage (Donlan, 2002; Donlan and Costerton, 2002; Stoodley *et al*., 2002; Hall-Stoodley *et al*., 2004). Formed in various environments the biofilms have different structures, depending on the type of the surface, physical conditions, in which they are formed, nutrients availability, and also genetic background and metabolic determinants of the bacteria themselves (Davey and O’Toole, 2000; Donlan, 2002; Ren *et al*., 2018; Sauer *et al*., 2022).

Studies on the process of biofilm formation have shown that certain cellular factors are at play when it comes to transition to sessile lifestyle. A crucial role is played by structures responsible for bacterial motility and also their efficient adhesion to surfaces, such as flagella, type IV pili or fimbriae (O’Toole and Kolter, 1998; Klausen *et al*., 2003). It has been demonstrated for *Pseudomonas aeruginosa* and *P. fluorescens* that non-flagellated mutants are defective in biofilm formation both on biotic and abiotic surfaces (O’Toole and Kolter, 1998; De Weger *et al*., 5275). But, some strains of *P. fluorescens* or *P. putida* do not require flagellum to attach to abiotic surfaces if certain sources of carbon (e.g. citrate) and iron ions are present in the environment (O’Toole and Kolter, 1998). The attachment abilities of bacteria, connected also with the features present on the cell membranes, are expressed in the synthesis of exopolysaccharides, including polysaccharides, alginate and lipopolysaccharides (LPS) (Nakao *et al*., 2012; Schurr, 2013). Absence of the O-antigen component of the LPS in *P. aeruginosa* and *P. fluorescens* prevented the bacteria to colonize abiotic surfaces and plant roots, respectively (Makin and Beveridge, 1996; Dekkers *et al*., 1998). However, it has also been shown that the type of carbon source (e.g. glucose or citrate) influences the pili-dependent aggregation (Tolker-Nielsen *et al*., 2000; Heydorn *et al*., 2002).

Bacterial cells embedded in biofilm matrix differ from the cells of the same bacterial species living as planktonic suspensions in the metabolic features they display, different growth rates and different expression and regulation of specific genes (Brown and Williams, 1985; Stoodley et al., 2002; Petrova and Sauer, 2012; MartÃnez and Vadyvaloo, 2014). The extracellular matrix formed by polymers (EPS, extracellular polymeric substances) composed of extracellular DNA (eDNA), proteins, and polysaccharides (Whitchurch *et al*., 2002; Friedman and Kolter, 2004; Ryder *et al*., 2007; Karatan and Watnick, 2009) protects the biofilm and serves as a structure enhancer. Protein synthesis and participation of ATP-dependent Clp proteases (*i.e*. ClpP) has also been shown to be required for biofilm formation in *P. fluorescens* (O’Toole and Kolter, 1998). Regulation of biofilm-related processes takes place at various levels in the bacterial cell, involving, among others, the highly conserved two-component regulatory system GacA/GacS (global activator of antibiotic and cyanide production), which also modulates the expression of genes involved in motility or response to stress factors, synthesis of antimicrobial compounds and siderophores (Haas and Keel, 2003; Raaijmakers *et al*., 2010; Cheng *et al*., 2013; Duque *et al*., 2013).

In case of plant-associated bacteria the formation of microcolonies on the host seeds, leaves or roots, is also considered as biofilm development, exhibited by pathogenic, mutualistic and neutral bacterial species (Bais *et al*., 2004; Timmusk *et al*., 2005; Fujishige *et al*., 2006; Ude *et al*., 2006; Rudrappa, Splaine, *et al*., 2008) and which may be the determinant of their competitive advantage in the environment (Davey and O’Toole, 2000). Numerous plant growth promoting rhizobacteria (PGPR) benefit from contact with host tissues and substances secreted by the plants, e.g. root exudates, in presence of which they engage the chemotactic mechanisms (Van de Broek *et al*., 1998). Studies undertaking the subject of bacterial biofilm development on biotic and abiotic surfaces show examples of correlation between biofilm formation on artificial structures and bacterial adhesion to surfaces of plant tissues, including root surface in the rhizosphere (Ramos *et al*., 2000; Yousef-Coronado *et al*., 2008; Fuqua, 2010). It was shown for *P. fluorescens* that a greater role in the colonization of roots is played by bacterial motility (Barahona *et al*., 2010). On the other hand, in case of *P. putida* it was demonstrated that mutations in genes encoding adhesins negatively influence both biofilm formation and bacterial attachment to biotic surfaces (Yousef-Coronado *et al*., 2008). Rhizosphere-associated strain of *P. putida* KT2440 was shown to attach to corn seeds through putative surface and membrane proteins (Espinosa-Urgel *et al*., 2000).

Much of the available data on biofilm formation and factors regulating it concern, however, mostly the well-described strains of *Pseudomonas aeruginosa*, *P. fluorescens* or *P. putida.* How the process is influenced by the environmental cues and what cellular components it involves in the little-known strains, such as the investigated in this work *P. donghuensis* P482, requires broader analysis. The P482 strain drew our attention since it exhibits not yet fully identified antimicrobial activity towards plant pathogenic bacteria and fungi, which is carbon source-dependent (Krzyzanowska et al., 2016; Ossowicki et al., 2017; Matuszewska et al., 2021; unpublished data). No typical quorum sensing systems, AHL-or quinolone-dependent, were identified in the P482 strain that would take part in communication between the P482 cells. The electron and atomic force microscopy observations demonstrated that the P482 cells are flagellated at one of the poles, what would enable the bacteria to move in the environment and possibly take part in cell attachment to various surfaces. Being a tomato rhizosphere isolate it was shown to colonize plant rhizosphere (Krzyzanowska *et al*., 2012) and respond adaptively to tomato and maize exudates (Krzyżanowska *et al*., 2023). Up to date, no data was available on the determinants of biofilm formation on biotic or abiotic surfaces by *P. donghuensis* P482 or any of its three sister strains (i.e. HYS^T^, SVBP6, P17) (Gao *et al*., 2012; Agaras *et al*., 2018; Marin-Bruzos *et al*., 2021), what prompted us to investigate the process from the point of bacterial cell and environment-derived factors. The influence of knock-outs in selected genes of P482, carbon source (glucose, glycerol or citrate), the type of colonized surface, abiotic (polystyrene or glass) or biotic, root tissues of tomato and maize, on biofilm-related characteristics of the strain was analysed.

## Experimental Procedures

### Bacterial strains and culture conditions

Bacterial strains used in this study were cultured in Miller’s Lysogeny Broth (LB) or on LB solidified with 1.5% agar (Novagen, Merck Group, Germany). When required, the media were supplemented with kanamycin (30 µg/ml) and/or gentamycin (40 µg/ml), for maintaining *P. donghuensis* P482 insertion mutants or GFP-tagged versions of the strains, respectively. For biofilm assays the strains were cultured in minimal M9 medium (Sambrook, 2001) supplemented with appropriate carbon source, as described in assays methodology. *P. donghuensis* P482 strains were grown at 28°C, *Escherichia coli* ST18 strain was grown at 37°C in media supplemented with 50 µg/ml of 5-aminolevulonic acid (5-ALA, Sigma-Aldrich, USA), with or without shaking (120 rpm shaking rate). For determination of bacterial growth rate the cells were cultured in minimal M9 medium supplemented with carbon source, in 96-well plates in 28°C with shaking and the OD_600_ measurements were performed hourly using an EnVision Multilabel Reader (Perkin Elmer, USA).

### Site-directed mutagenesis

The P482 mutants analysed in this study were obtained as previously described in (Krzyzanowska *et al*., 2016). Briefly, fragments (392–624 bp) of genes selected for site-directed mutagenesis were amplified and cloned in the XbaI/XhoI, XbaI/KpnI or XbaI/ApaI restriction sites of the pKNOCK-Km suicide vector (Alexeyev, 1999). The resulting constructs named pKN0543, pKN0864, pKN0868, pKN0887, pKN0892, pKN0894, pKN0850, pKN0883, pKN1101, pKN1102, pKN1251, pKN1850, pKN3824, pKN4694, pKN4697, pKN4698, pKN4700, pKN5088, pKN5092, pKN5095 were introduced into the *E.coli* ST18 donor strain (Thoma and Schobert, 2009) and subsequently transferred to *Pseudomonas* sp. P482 by biparental mating. The P482 transconjugants were screened for the presence of the pKNOCK-Km insertion with primers F_pKNOCK_backbone and R_pKNOCK_backbone (see Matuszewska et al., 2021 for details). The site of incorporation of the suicide vector into the target loci was confirmed through a sequencing reaction using the primer F_outof_pKNOCK (Matuszewska *et al*., 2021) with genomic DNA of each mutant as a template. This allowed for mapping of the pKNOCK-Km insertion site to the genome of the P482 strain. The sequencing was performed at Oligo.pl (Warsaw, Poland). Sequences of the used primer pairs, annealing temperatures and the expected lengths of amplicons are given in Supplementary Table S1.

### Fluorescent labelling of P482 strains

GFP-expressing derivatives of the *P. donghuensis* P482 strains were obtained by transforming the bacteria cells with pPROBE-GTkan vector (Miller *et al*., 2000) by electroporation (Gene Pulser Xcell; Bio-Rad, USA), as described in (Krzyzanowska *et al*., 2012), with default pulse conditions set for the *Pseudomonas* strains. The transformants were selected by plating the bacterial suspensions post-electroporation on LB-agar plates supplemented with kanamycin and gentamicin.

### Motility assays

Swimming motility was determined as the diameter of zone travelled by bacteria inoculated with a pipette tip into 0.3% agar M8 plates, supplemented with 0.02% glucose, 0.05% casamino acids and 1 mM MgSO_4_, and incubated for 18–20 h at 28°C (Ha et al., 2014, modified). Swimming diameter was measured post-incubation using ImageJ. The data was analysed using GraphPad Prism.

Swarming motility was assayed by applying bacteria on the surface of 0.8% agar M8 plates supplemented with 0.1% glucose and 0.25% casamino acids and 1 mM MgSO_4_ (Ha et al., 2014, modified). Swarming plates were incubated for 18-20 h at 28°C. Swarming diameter was measured post-incubation using ImageJ. The data was analysed using GraphPad Prism 9.5.1 (GraphPad Software, USA).

Twitching motility was analysed on 1.5% agar M63 plates supplemented with 0.02% glucose, 0.05% casamino acids and 1 mM MgSO_4_ (Ha et al., 2014, modified). Twitching plates were incubated for 48 h at 28°C and the results were analysed qualitatively post-incubation.

Bacterial motility in liquid medium was determined using the motility medium S Base (HiMedia, USA), prepared according to the manufacturer’s protocol. The medium was inoculated using inoculation loop with overnight bacterial cultures from LA plates. The tubes were incubated at 28°C without shaking for 24 hours, after that the bacteria growth was observed and photographed. The assay was performed in three replicas.

### Colony biofilm assay

The colony morphology was assessed in Congo red assay as described in (Wang *et al*., 2021). Briefly, plates with M63 agar supplemented with 0.2% glucose, 0.5% casamino acids, 1 mM MgSO_4_, 1.5% agar, and 2 ml of a 50x Congo red-Coomassie blue solution (2 mg/ml Congo red and 1 mg/ml Coomassie blue) per 100 ml of medium were prepared. Three microliters of overnight cultures of each strain were applied on the Congo red plates, which were incubated at 28°C for 24 hours and then left at room temperature for 5 to 7 days for biofilm formation. Images of the colonies’ morphology were taken with Leica MZ10F stereomicroscope (Leica, Germany).

### Biofilm formation assays

#### Crystal violet staining

The assay analysed the ability of the bacteria to form biofilm on the polystyrene surface in 96-well microtiter plates (O’Toole, 2010, with modifications). Overnight cultures of the analysed strains were diluted 1:100 in minimal M9 medium supplemented with 0.4% glucose, 0.4% glycerol, or 20 mM sodium citrate. 150 µl aliquots were dispensed to wells of the microtiter plates (Sarstedt, Germany), in 4 replicates each and incubated overnight at 28°C, without shaking. Following incubation the plates were washed twice with distilled water to remove planktonic cells. The attached biofilm was then stained using 150 µl of 0.1% crystal violet solution for 20 minutes, and washed three times with distilled water to remove the excess stain. The biofilm-bound dye was re-solubilized by adding 150 µl of 30% acetic acid to each well. Absorbance at 550 nm was measured after 20 min incubation at room temperature, with the use of Envision Multilabel plate reader (Perkin Elmer, USA). Biofilm formation was calculated by normalizing for bacterial growth using OD_550/600_. The results obtained for mutant strains were normalized to 100% taken as the absorbance value measured for the wild-type P482 strain. Analysis of variance, ANOVA, was performed for each of the analysed P482 strains with multiple comparisons and Dunnett’s or Tukey test used to correct for multiple comparisons, to determine differences between and within strains. The statistical analysis was done using GraphPad Prism 9.5.1 (GraphPad Software, USA).

#### Biofilm formation on glass

Biofilm formation on glass surface was assessed in static conditions, as follows: overnight cultures of GFP-tagged *P. donghuensis* P482 strains were diluted 1:500 into fresh M9 minimal medium supplemented with appropriate carbon source (0.4% glucose, 0.4% glycerol, or 20 mM sodium citrate), in glass-bottom 24-well microtiter plates (SensoPlate^TM^, Greiner Bio-One, Austria). The bacteria were incubated for 24 h at 28°C without shaking; prior to imaging the medium containing non-attached cells was removed and the biofilms were covered with phosphate-buffered saline (PBS). Plates were placed under a scanning confocal microscope Leica SP8X (Leica Microsystems, Germany), using a 10x objective. Z-stack images were collected for every well, with average step height of 3.85 µm; the limits of stacks were set by moving within the z axis range, until no fluorescence was observed for both upper and lower stack. The images were acquired and processed via Leica Application Suite X (LAS X, Leica, Germany) software, with four randomly selected fields of each sample evaluated. The 2D images were constructed from 3D z-stack images using the Maximum Intensity Projection algorithm in LAS X by selecting pixels of the highest intensity from every slice. Fluorescence data from the collected images were obtained by measuring the mean fluorescence intensity using the ImageJ software. Analysis of variance, ANOVA, was performed for each of the analysed P482 strains with multiple comparisons and Dunnett’s or Tukey test used to correct for multiple comparisons, to determine differences between and within strains. The statistical analysis was done using GraphPad Prism 9.5.1 (GraphPad Software, USA).

### Tomato rhizosphere colonization

Tomato seeds (*Solanum lycopersicum L.*, *cv*. Saint Pierre, Vilmorin Garden, Poland) were surface sterilized by placing 20-30 seeds in a sterile Eppendorf tube and washing them 3-4 times with sterile distilled water, then for 1 minute in 70% ethanol, 3 minutes in 3% NaOCl, and again 3-4 times with sterile water. The seeds were placed on 0.5x Murashige-Skoog agar plates and left for germination in dark for 4 days at 25°C, to select for those not contaminated with microorganisms. Subsequently, the seedlings were inoculated with suspensions of the GFP-tagged versions of the analysed P482 strains in 10 mM MgCl_2_ at OD_McFarland_=6. For every P482 strain three germinated seeds were placed in a Petri dish, each covered with 10 µl of respective bacterial suspension, and incubated at room temperature for 20 minutes. The inoculated seedlings were placed in Magenta boxes (Sigma-Aldrich, USA) containing 100 g sterile grit and 11 ml 1x Murashige-Skoog medium supplemented with 3% sucrose. Seedlings covered with 10 mM MgCl_2_ only served as negative control. The boxes were incubated in a phytotron chamber for 4 weeks (16/8 h photoperiod, 20°C). After incubation, root samples were prepared for microscopy, as described in (Krzyzanowska *et al*., 2012) with modifications. Briefly, roots were cut and placed in Petri dishes (6 cm diameter; Sarstedt, Germany) and enrichment medium (2 g/l KH_2_PO_4_, 4.66 g/l K_2_HPO_4_, 1.33 g/l (NH_4_)_2_SO_4_, 50 mg/l FeSO_4_, 80 mg/l MgSO_4_, 10 g/l agar, 0.4% glycerol, 40 mg/l gentamicin) was poured over the samples. The samples were incubated overnight at 28°C and analysed with a Leica SP8X Confocal (Leica Microsystems, Germany), using a 10x objective and GFP excitation/detection settings. Multiple images from different parts of roots of the analysed samples were taken for evaluation of tissue colonization by bacteria. The entire experimental setup was repeated two to four times for each analysed P482 derivative.

### Maize rhizosphere colonization

Maize seeds (*Zea mays* L., *cv*. Bejm, Plant Breeding and Acclimatization Institute, Smolice, Poland) were surface sterilized by immersion in 5% NaOCl for 15 min, in 70% ethanol for 10 min, and subsequent wash with sterile distilled water. The seeds were placed on water agar plates (15 g/l) and incubated in phytotron chamber for 7 days for sprouting (16/8 h photoperiod, 20°C). For inoculation of the maize sprouts, saline suspensions were prepared from the overnight cultures of GFP-tagged *P. donghuensis* P482 wild-type and derivative strains, OD_McFarland_ =1. Maize sprouts were immersed in the bacterial suspensions, two per strain, in Petri dishes and incubated for 30 min at room temperature. Next, the sprouts were placed in sealed conical flasks containing 20 ml of sterile water and placed in the phytotron chamber for further 7 days for growth. Maize sprouts immersed in clear saline served as negative control. Preparation of root samples and enrichment procedure followed as described for tomato colonization, with the root samples cut and placed in square Petri dishes (10×10 cm; VWR, USA) and the enrichment medium was poured over them. The samples were incubated overnight at 28°C and analysed with a Leica HCS LSI Macro Confocal (Leica Microsystems, Germany), using a 5x objective and GFP excitation/detection settings. Multiple images from different parts of roots of the analysed samples were taken for evaluation of tissue colonization by bacteria. The entire experimental setup was repeated at least twice for each analysed P482 derivative.

## Results

### Motility of the *P. donghuensis* P482 cells is mostly dependent on the activity of flagella

To analyse cellular factors involved in the process of biofilm formation by the P482 strain a genome analysis was performed and a twenty knock-out mutants were constructed by means of site-directed mutagenesis using a suicide pKNOCK vector (Alexeyev, 1999). The target genes of the obtained mutants were grouped into three subgroups (see Table 1) as those involved in motility or chemotaxis (Belas, 2014; Sampedro *et al*., 2015; Bouteiller *et al*., 2021), genes encoding proteases and the *gacA* regulatory gene (O’Toole and Kolter, 1998; Heeb and Haas, 2001), and genes engaged in synthesis of components of the biofilm matrix and adhesion (Nakao *et al*., 2012; Schurr, 2013). The genome of *P. donghuensis* P482 contains almost forty genes encoding proteins which have a potential to build the motility organelle. The knock-outs included the *flgL* gene (mutant KN0864) encoding the flagellar hook-associated protein, *fliC* encoding flagellin, being the main component of the filament (KN0868), *fliM* (KN0887) – coding for the motor switch protein which allows for motion and rotation of the flagellum, and *fliR* and *flhA* (KN0892 and KN0894, respectively) encoding proteins involved in flagellar export.

**Table 1.** Selected genes encoding proteins possibly involved in biofilm formation by *P. donghuensis* P482, targeted by site-directed mutagenesis.

| Mutant's name | Locus | Annotated gene | Product | Function in biofilm formation |
| --- | --- | --- | --- | --- |
| KN0864 | BV82_0864 | *flgL* | flagellar hook-associated protein 3 | Motility/Chemotaxis |
| KN0868 | BV82_0868 | *fliC* | bacterial flagellin N-terminal helical region family protein |  |
| KN0887 | BV82_0887 | *fliM* | flagellar motor switch protein FliM |  |
| KN0892 | BV82_0892 | *fliR* | flagellar biosynthetic protein FliR |  |
| KN0894 | BV82_0894 | *flhA* | flagellar biosynthesis protein FlhA |  |
| KN0850 | BV82_0850 | *cheW* | chemotaxis protein CheW |  |
| KN0883 | BV82_0883 | *cheY* | chemotaxis protein CheY |  |
| KN1101 | BV82_1101 | *clpP* | ATP-dependent Clp protease proteolytic subunit ClpP | Proteases/Regulatory |
| KN1102 | BV82_1102 | *clpX* | ATP-dependent Clp protease ATP-binding subunit ClpX |  |
| KN1251 | BV82_1251 | *clpA* | ATP-dependent Clp protease ATP-binding subunit ClpA |  |
| KN3318 | BV82_3318 | *gacA* | GacA, a response regulator of the two-component GacS/GacA system |  |
| KN4694 | BV82_4694 | *algE* | alginate production protein AlgE | Biofilm matrix/Adhesion |
| KN4697 | BV82_4697 | *algL* | alginate lyase |  |
| KN4698 | BV82_4698 | - | alginate O-acetyltransferase |  |
| KN4700 | BV82_4700 | *algF* | alginate O-acetyltransferase AlgF |  |
| KN5088 | BV82_5088 | - | polysaccharide biosynthesis family protein |  |
| KN5092 | BV82_5092 | - | O-antigen ligase family protein |  |
| KN5095 | BV82_5095 | - | polysaccharide biosynthesis/export family protein |  |
| KN1850 | BV82_1850 | - | lipopolysaccharide kinase family protein |  |
| KN0543 | BV82_0543 | - | lipid-A-disaccharide synthase |  |
| KN3824 | BV82_3824 | - | adhesin |  |

The flagellum synthesis mutants were analysed together with all other mutants (listed in Table 1) for their swimming, swarming and twitching motilities on appropriate agar-solidified media and in semi-liquid motility medium S. Swimming motility was determined with a plate assay on 0.3% agar-solidified M63 medium (Rashid and Kornberg). All five flagellum synthesis-related mutants (KN0864, KN0868, KN0887, KN0892 and KN0894) showed defects in swimming motility when compared to the wild-type P482 strain (Fig. 1A and B). The bacteria were not able to move further from the stabbing point outwards to the margins of the plate. Similar result was obtained in the assay performed in the liquid medium S (Fig. 1C) – none of the mutants with disturbed flagellum protein synthesis was capable to move away from the inoculation line. The remaining mutants analysed in the two assays showed no significant changes in the size of the diameter of swimming halo or spreading in the liquid medium when compared to the wild-type strain (see Supplementary Fig. S1A-C). This shows that the swimming motility of the P482 cells is flagellum-dependent.

**Figure 1.**
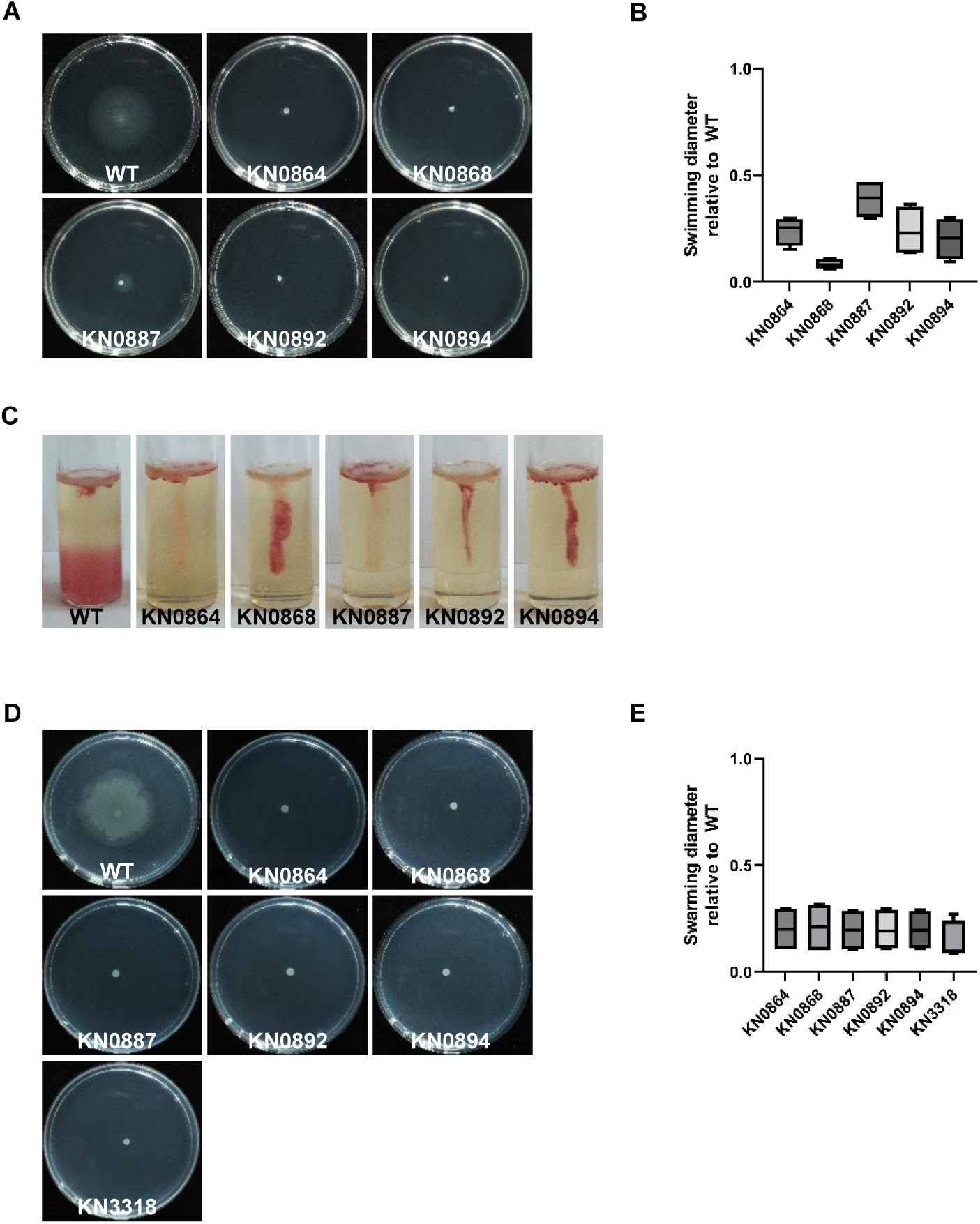
Motility phenotypes of the P482 strains. (A) Images of P482 mutants demonstrating swimming defect versus the wild-type strain. (B) Swim diameters of the mutant strains relative to WT (taken as 1), n=4. (C) Results of a motility assay in motility medium S showing non-motile P482 mutants. (D) Images of P482 mutants demonstrating swarming defect versus the WT strain. (D) Swarm diameters of the mutant strains relative to the WT (taken as 1), n=4 or 5 replicates.

The swarming motility which allows the bacterial cells to propagate on semi-solid surface (Köhler *et al*., 2000) was assessed for the P482 mutants on 0.8% agar plates. Similarly as in the swimming motility analysis, the five mutants in flagellum-related genes were entirely unable to swarm (Fig. 1D and E), as compared to the wild-type P482 which demonstrated swarming ability in a “bull’s eye” or terrace type pattern of the halo (Kearns, 2010) and the growth of the mutants was visible only where the culture was applied. The only other mutant which also demonstrated defect in the swarming motility was the *gacA* mutant (KN3318). One mutant, KN0850, with knock-out in the *cheW* chemotaxis protein encoding gene, occurred to be hypermotile in the swarming test (Supplementary Fig. S2A and B). These results suggest that swarming in *P. donghuensis* also depends mostly on the presence of efficient flagella, however, the activity of the *gacA* response regulator gene/other factors seems to play an important role too (Kinscherf and Willis, 1999).

For the twitching analysis, dependent on the activity of type IV pili where cells move across solid surfaces (Bertrand *et al*., 2010) it was the most challenging to adjust the experimental conditions to actually observe this type of motility in P482 cells. Since the satisfying results were obtained only for one repetition of the assay, they will be analysed qualitatively with cautiousness. In only two of the analysed mutants it was possible to observe defect of twitching activity exhibited in the lack of halo of colony expansion by cells moving sub-surface of the used medium (Supplementary Fig. S3). The mutants were KN4698 and KN0543, in genes encoding alginate O-acetyltransferase and lipid-A-disaccharide synthase, respectively. Mutants in genes connected to synthesis of flagellum which were affected in swimming and swarming motilities showed twitching pattern similar to that of the wild-type strain, which shows that activity of flagellum is not essential for this type of movement in P482. Overall, the results suggest that bacterial motility of *P. donghuensis* P482 is connected to the active, mobile flagella in case of swimming and swarming motilities but twitching seems to depend on other factors, not involving the presence of this organelle, and what might have implications also for the ability of the P482 cells to use different modes of attachment to surfaces.

### Biosynthesis of biofilm matrix components involves activity of distinct cellular factors

The synthesis of various polymers composing the matrix of biofilm, i.e. exopolysaccharides or lipopolysaccharides, has been shown to be crucial for cells auto-aggregation and biofilm formation in various bacteria of the *Pseudomonas* genus, including plant-associated strains (Ryder *et al*., 2007; Ghafoor *et al*., 2011; Bogino *et al*., 2013; Maunders and Welch, 2017). One method which allows to determine the ability of bacterial cells to produce EPS is Congo red binding assay (Friedman and Kolter, 2004; Wang *et al*., 2021). The Congo red is a diazo dye that binds to amyloid fibers and biofilm matrix polysaccharides (Jones and Wozniak, 2017). In the assay the morphology of bacterial colonies is assessed qualitatively, and differences in the distribution of the bound dye and colour intensity can be observed. The colony of the wild-type strain of P482 shows light pink pigmentation with the colour evenly distributed in the bacterial mass (Fig. 2), what might suggest relatively low levels of EPS production by the strain in this experimental setup (Wang *et al*., 2021). Nevertheless, two of the analysed mutants showed no (KN3318 – *gacA*) or much reduced (KN1850) Congo red binding. The colony of the *gacA* mutant was lacking any red pigment, possibly pointing to regulation of the synthesis of the extracellular polymeric substances in P482 by the *gacA/gacS* system (Wei and Ma, 2013; Li *et al*., 2015). The KN1850 mutant with a slightly lighter colour of the colony is a knock-out in a gene encoding a lipopolysaccharide kinase family protein, what suggests the loss of its activity resulting in defects in LPS production (Bertani and Ruiz, 2018). An interesting group of mutants are those where the colonies show uneven distribution of the red pigment, with darker red sections in the centre of the colony or dark-red stripes in the peripheral parts. The first is the case of colonies formed by KN0883 and KN0887, mutants in *cheY* gene encoding chemotaxis protein CheY and *fliM* encoding FliM protein being the flagellar motor switch, respectively. A ‘striped’ colony phenotype can be observed for two mutants, KN1101 and KN1102, in genes encoding ClpP and ClpX proteases, respectively; for strains with knock-outs in genes encoding O-antigen ligase family protein (KN5092), and polysaccharide biosynthesis/export family protein (KN5095), what might mean local accumulation of matrix polymers as result of disturbed activity of the enzymes involved in their synthesis and/or transport (Figaj *et al*., 2019). For two mutants, KN4700, where *algF* gene encoding alginate O-acetyl transferase AlgF was inactivated, and KN0543, with lipid-A-disaccharide synthase encoding gene not active, a darker red colour of the colony, with more dye bound to the matrix, was observed, hinting at higher levels of polysaccharides’ synthesis in those mutants possibly utilizing pathways which do not involve activity of the two enzymes.

**Figure 2.**
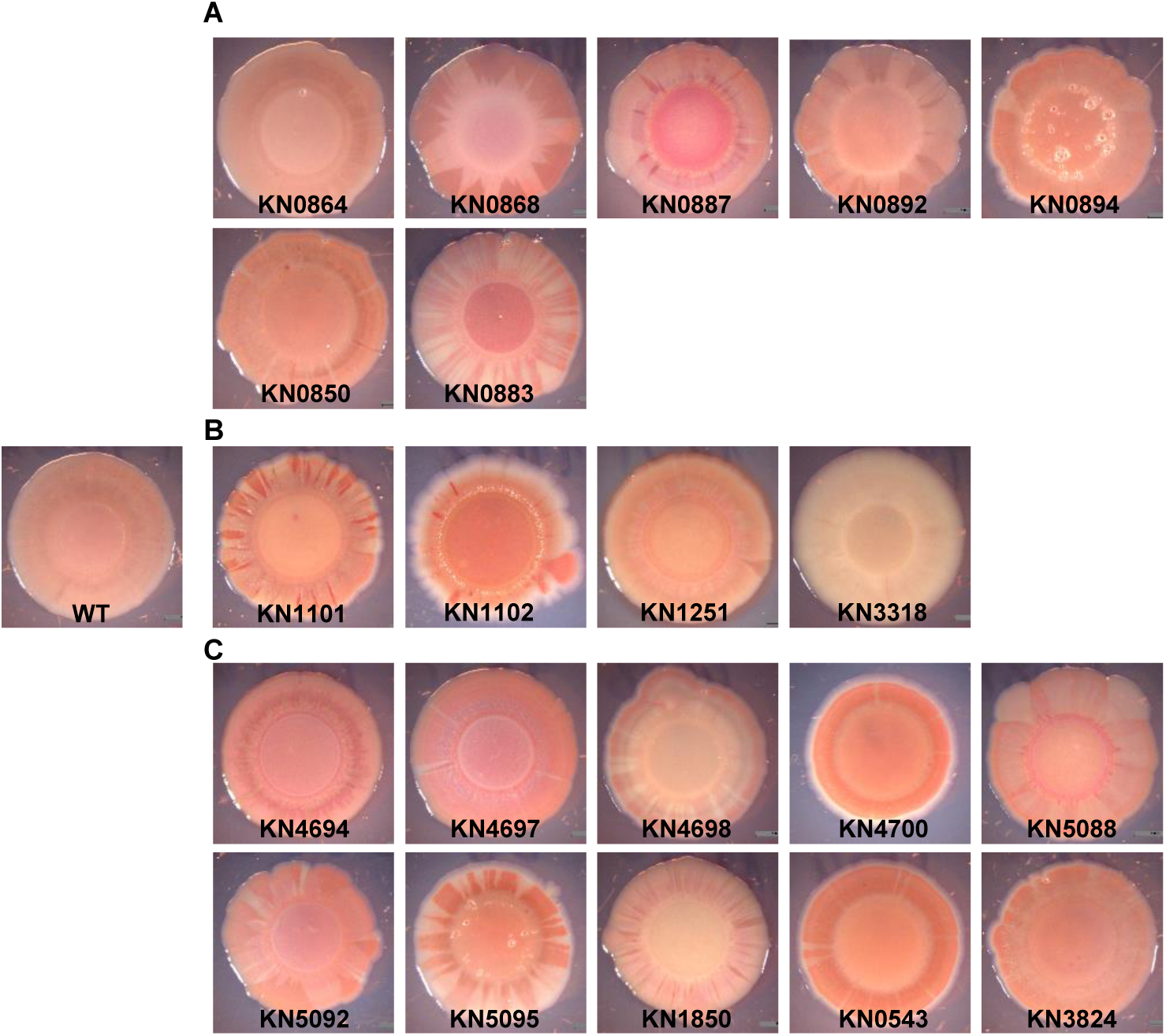
Biofilm phenotypes – colony morphology of the P482 wild type and mutant strains. Images of Congo-red stained colonies of the P482 WT and mutant strains: (B) knock-outs in motility and attachment-related genes, (C) – knock-outs in proteases-encoding and *gacA* (KN3318) regulatory genes, (D) – knock-outs in matrix synthesis-related genes.

### Biofilm formation by *P. donghuensis* P482 is affected by carbon source

Transition from planktonic to sessile lifestyle and formation of biofilm by bacteria is directly affected by the availability of nutrients in the environment (Davey and O’Toole, 2000; Rossi *et al*., 2022) of which carbon sources play possibly one of the leading roles (Sauer *et al*., 2004; Harmsen *et al*., 2010; Huynh *et al*., 2012). Great deal of data on the regulation of biofilm formation is, however, available for *P. aeruginosa* mostly, therefore it was interesting to ask how carbon sources might affect this activity in a less-studied environmental strain which is *P. donghuensis* P482. A biofilm formation assay was performed in a 96-well microplate setup (O’Toole, 2010) where the wild-type and mutant strains were cultured in M9 minimal medium supplemented either with glucose, glycerol or citrate and the biofilm formed in the wells (at the bottom and the air-liquid interface) was stained with crystal violet. The dye binds to the cells attached to a surface, what, after re-solubilization of the dye with acetic acid, allows for quantitative analysis of the formed biomass. The defined minimal mineral medium was chosen for this analysis to enable comparison of effects of carbon source on biofilm formation in a direct way, what is not quite possible when complex media, such as standard LB (Luria broth) or TSB (tryptic soy broth), are used.

First, the wild-type P482 strain was analysed for the effect of a specific carbon source on the biofilm formation process. The absorbance measurements done post-staining were normalized to OD_600_ of the bacterial cultures before staining. Differences in biofilm formation in the three carbon sources were observed: the biofilm was formed most efficiently when glycerol was present in the medium (Fig. 3A), with glucose the production of biofilm was ca. two fold lower, and the presence of citrate resulted in the least efficient formation of biofilm. This showed that the carbon source is an important factor in the process of biofilm formation by the wild-type *P. donghuensis* P482. From this point it was only natural to verify how glucose, glycerol and citrate affect the biofilm in the P482 mutants. None of the mutants showed decrease in biofilm production in the presence of glucose (Fig. 3B-D); three mutants in the motility-related genes, namely KN0887 (*fliM*), KN0892 (*fliR*) and KN0894 (*flhA*) showed significant increase (up to and over 200%) in biofilm formation when compared to the wild-type strain (Fig. 3B), suggesting that in this experimental setup glucose favors transition of non-motile cells to surface-attached. Similar effect was observed for the mutant in one of the protease genes, KN1102 (*clpX*), and in the regulatory gene *gacA* (KN3318), with over two-and over three-fold increase in biofilm formation, respectively (Fig. 3C). Also, two mutants KN4694 and KN4698, both involved in alginate synthesis in the cell, formed twice as much biofilm as the wild-type when glucose was present in the medium (Fig. 3D).

**Figure 3.**
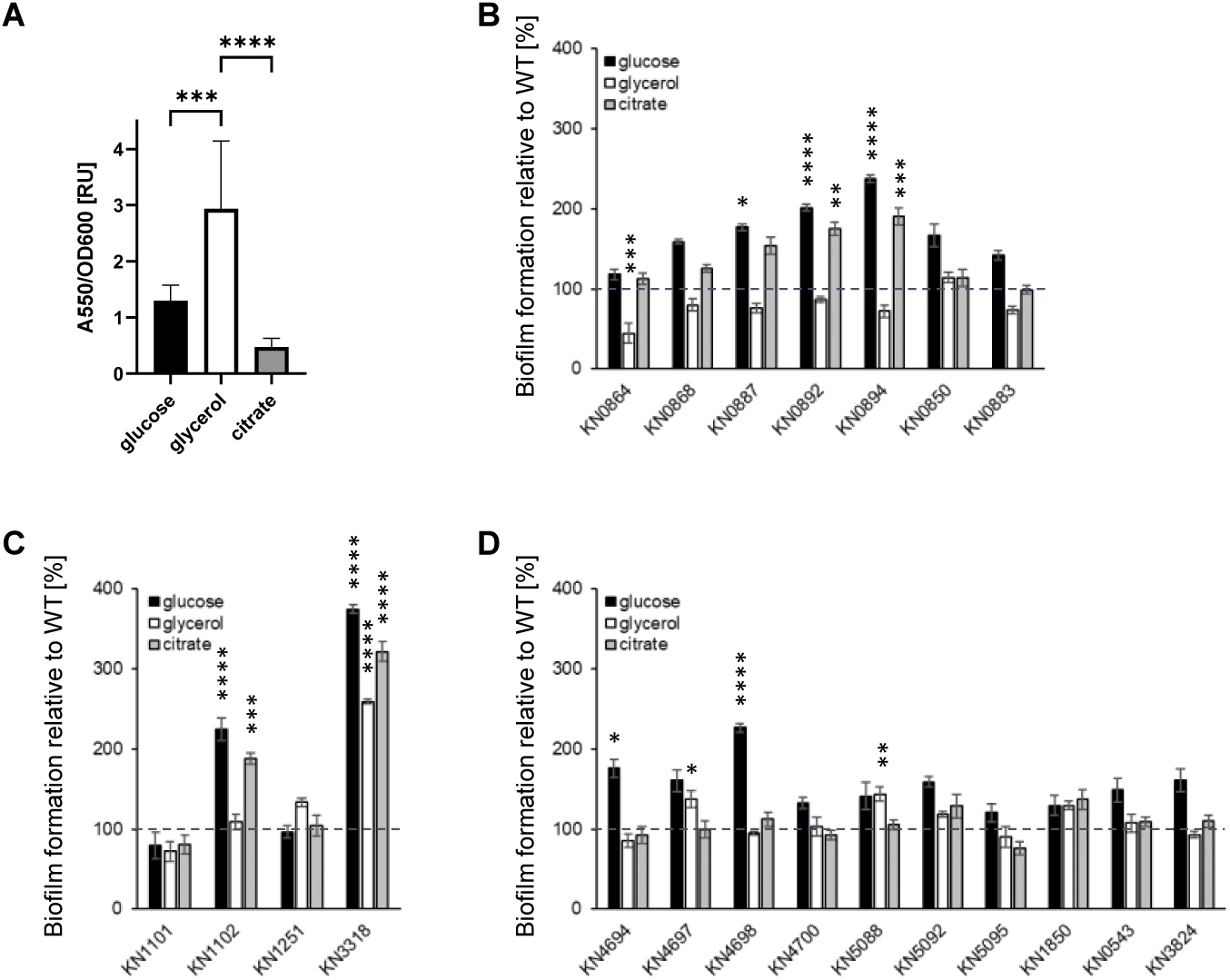
Biofilm formation by P482 mutant strains in crystal violet binding assay. Biofilm formation by P482 WT (A) and mutant strains (B) – knock-outs in motility and attachment-related genes, (C) – knock-outs in proteases-encoding and *gacA* (KN3318) regulatory genes, (D) – knock-outs in matrix synthesis-related genes, on abiotic surface (polystyrene) in M9 minimal medium in relation to provided carbon source (0.4% glucose, 0.4% glycerol, 20 mM citrate). Biofilm formation by the mutant strains was determined by absorbance measurements normalized to those obtained for WT, taken as 100%. Grey dashed line marks 100% of biofilm formation by the WT strain. See Materials and Methods section for details of the procedure. Absorbance was measured at 550 nm. The results represent mean values for three independent experiments with three sample replicates each. The error bars represent SEM. Statistical significance was determined with ANOVA and Dunnet’s multiple comparisons test for correction (*P < 0.05; **P < 0.01; ***P < 0.001; ****P < 0.0001); all P values for the ANOVA analyses can be found in Supplementary Table S2 and S3.

When the minimal medium was supplemented with glycerol most of the mutants formed biofilm similar to that of the wild-type strain, suggesting that the mutations did not affect cells’ attachment when this carbon source was available. Only one mutant, KN0864, in the *flgL* gene encoding flagellar hook-associated protein, demonstrated over 50% decrease in biofilm formation (Fig. 3B). Three mutants showed increased ability to form biofilm in glycerol: KN3318, knock-out in the *gacA* gene, which similarly as in glucose showed nearly 2.5 times higher efficiency in forming biofilm than the wild-type; and two mutants in genes associated with polysaccharide degradation and synthesis, KN4697 (gene encoding alginate lyase) and KN5088 (gene encoding polysaccharide biosynthesis protein) (Fig. 3D). What needs to be taken into account is that in glycerol *P. donghuensis* P482 strains, both wild-type and mutants, exhibit much slower growth and demonstrate a long lag phase (see Supplementary Fig. S3).

In case of citrate being available as a carbon source four mutants showed increase in biofilm formation: KN0892 (*fliR*) and KN0894 (*flhA*), both produced biofilm nearly twice as efficiently as the wild-type P482 (Fig. 3B); the other two mutants were KN1102 and KN3318, knock-outs in *clpX* and *gacA* genes, respectively, with the latter producing three times more biofilm than wild-type (Fig. 3C). For other analysed mutants the presence of citrate caused no significant differences in biofilm formation when compared to the wild-type strain. Overall, the results of the analyses show that the formation of biofilm on abiotic surface such as polystyrene in *P. donghuensis* P482 is a complex process, which is regulated on many levels and its efficiency depends on certain carbon sources available for bacteria.

### The type of abiotic surface has an impact on biomass production and biofilm structure in *P. donghuensis* P482

The effect of carbon source on biofilm formation observed in the crystal violet assay brought information on the differences in relative biofilm biomass accumulated by P482 strains. What was not determined was whether the carbon sources available for the strains during the process of biofilm formation by P482 have an impact on its structure. For the purpose of the structural analyses of biofilm with the use of confocal laser scanning microscopy (CLSM), the P482 strains, wild-type and mutants, were fluorescently tagged with the pPROBE-GT-Kan plasmid, expressing green fluorescent protein (GFP). The strains were cultured in glass-bottom 24-well plates, in minimal M9 medium supplemented with glucose, glycerol or citrate, in static conditions, what allows for microscopy observations of the formed biofilm structures. The CLSM images was obtained with z-stacks collected every 3.85 µm from the bottom to the top of the formed structures of the biofilm, with their number depending on the thickness of the biofilm. For details of the data analysis see details in Experimental procedures section.

The images obtained for the wild-type strain in the presence of the three different carbon sources revealed significant differences in the structure of the biofilm formed by the P482 strain. In glucose-supplemented medium the bacteria formed relatively thin layer, with single microcolonies of aggregating cells scattered on the surface (Fig. 4A and B). For glycerol the structure of biofilm was much more developed, the cell layer was thicker, with multiple microcolonies attached to each other, forming three-dimensional structures covering nearly the entire surface of the glass. In medium where citrate was the carbon source the biofilm formed by the wild-type P482 exhibited intermediate phenotype when compared to that in glucose and glycerol – the cells were clustered into more microcolonies than it was observed for glucose, but the biofilm was not as dense and mature as it was in case of glycerol (Fig. 4A and B). The effects observed *via* confocal microscopy confirmed glycerol as the carbon source promoting biofilm formation by *P. donghuensis* P482, when compared with glucose and citrate (Fig. 4C). However, biofilm formed on glass in presence of citrate was more developed than the one in glucose, which was opposite to the result obtained for polystyrene and crystal violet staining assay. One might look for the reason for this in the different nature of the abiotic surfaces used in the assays and their physicochemical properties.

**Figure 4.**
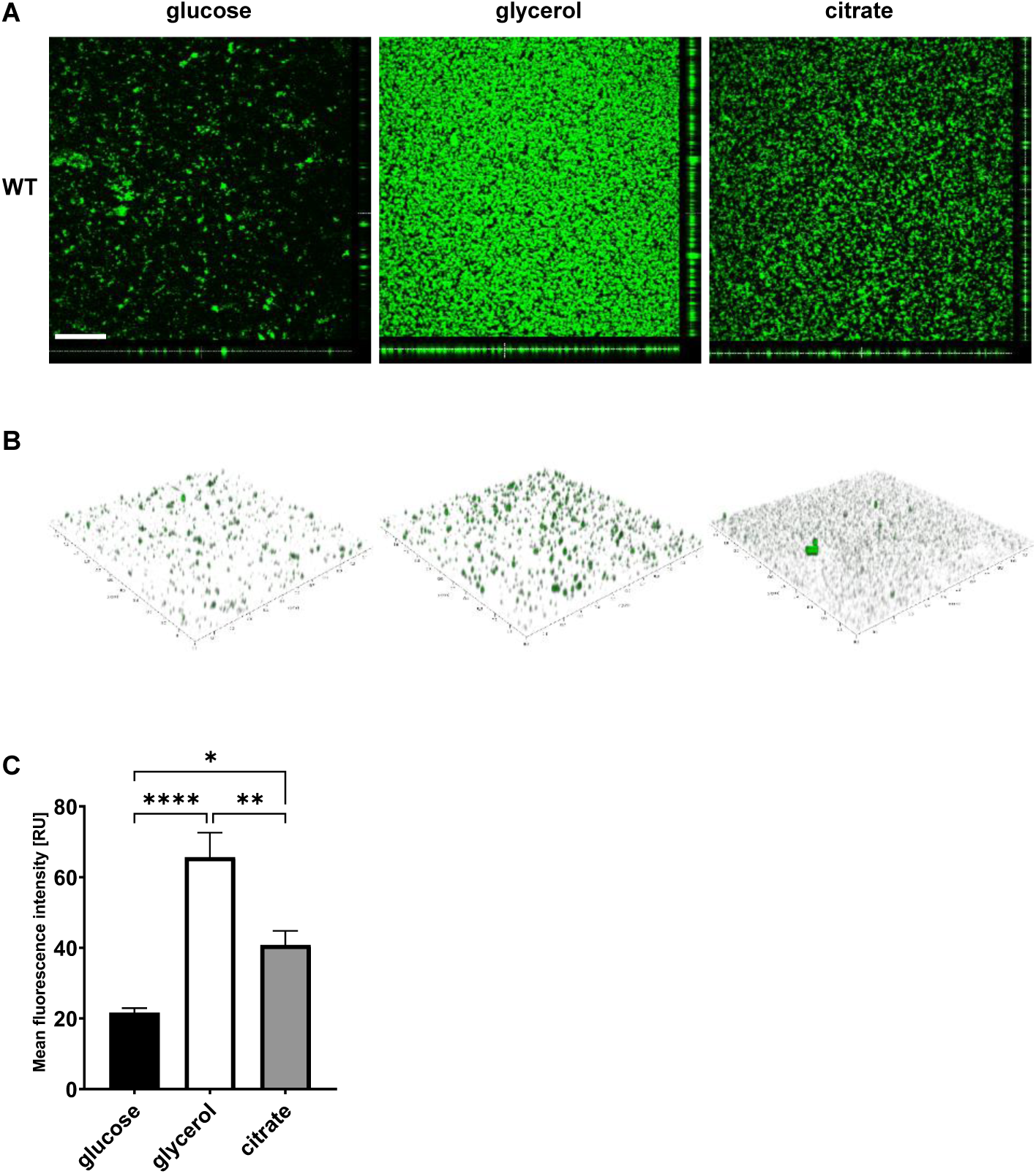
Biofilm formation by P482 wild type strain on glass, in relation to carbon source. (A) Orthogonal and (B) 3D view of biofilm formed by GFP-tagged WT P482 strain on the glass bottom of 24-well plates in M9 minimal medium supplemented with 0.4% glucose, 0.4% glycerol or 20 mM citrate. (C) Quantification of mean fluorescence intensity for the biofilm formed by the GFP P482 in the M9 medium with the given carbon sources. The bars represent means with SEM. Statistical significance was determined by one-way ANOVA (*P < 0.05; **P < 0.01: ****P < 0.0001). Bar in (A) = 200 µm.

The effect of glass surface on the P482 biofilm structure was observed in all three groups of analysed mutants. All mutants with knock-outs in flagellum-synthesis genes (KN0864, KN0868, KN0887, KN0982 and KN0894) exhibited a diffused biofilm phenotype, regardless of the carbon source present in the medium (Fig. 5A). Neither in glucose nor glycerol or citrate any of the five strains formed clustered microcolonies of attached cells, with the biggest differences observed for glycerol when compared to the wild-type strain. The cells, however, did form some sort of biofilm which did not detach easily from the surface during washing. When taking into consideration the mean fluorescence reads and calculations showing indirectly the amounts of cells engaged in formation of the layers, the biomass of the biofilm was still considerable in case of mutants KN0864, KN0868 and KN0894 when glucose was the carbon source (Fig. 5B). This suggests that in the presence of glucose it might not be the flagella that determine the initial attachment to the surface but they might be important for cells’ aggregation into microcolonies. In glycerol and citrate, on the other hand, the KN0864, KN0868, KN0887 and KN0883 mutants (the latter in *cheY* gene, encoding chemotaxis protein) demonstrated significant decrease in biofilm-forming abilities, exhibited in single layers of cells attached to the surface. Interesting result was obtained for the KN0850 mutant, knock-out in chemotaxis protein CheW-encoding gene, which in glucose-supplemented medium formed three-dimensional biofilm with much better developed structure than the wild-type strain did. In glycerol it formed thinner biofilm but still with distinguishable microcolonies, and a thin layer of cells with single microcolonies in the presence of citrate.

**Figure 5.**
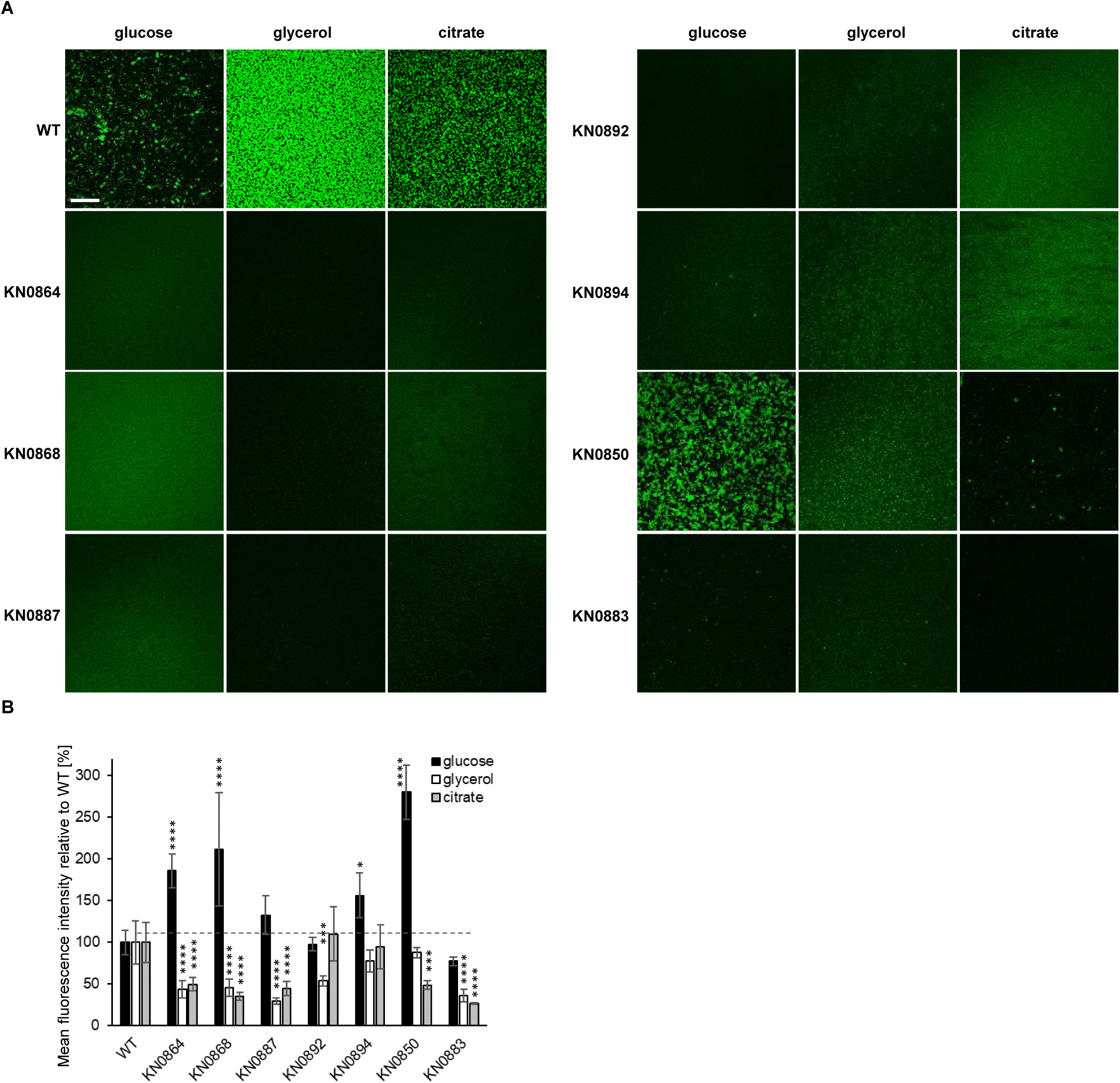
Biofilm formation on glass by GFP-tagged P482 mutants in motility and attachment-related genes. (A) Representative CLSM images of biofilm formed by GFP-tagged P482 strains on the glass bottom of 24-well plates in M9 minimal medium supplemented with 0.4% glucose, 0.4% glycerol or 20 mM citrate. Bar = 200 µm. (B) Quantification of mean fluorescence intensity for the biofilm formed by the GFP P482 mutants in the M9 medium supplemented with the individual carbon sources. Statistical significance for mutants vs. WT was determined with ANOVA and Dunnet’s multiple comparisons test for correction (*P < 0.05; **P < 0.01; ***P < 0.001; ****P < 0.0001); all P values for the ANOVA analyses can be found in Supplementary Tables S2 and S3.

In case of mutants with knock-outs in genes encoding proteases, ClpP and ClpX (KN1101 and KN1102, respectively), the effect observed for all three tested carbon sources was lack of structured biofilm where cell would form microcolonies (Fig. 6A) with a significant decrease of biomass production in the presence of glycerol and citrate (Fig. 6B). Glucose did not have that effect when it comes to biomass produced by the protease mutants – the cells accumulated into biomass at the surface of the glass but in a diffused manner (Fig. 6A and B). This suggests that activity of ClpP and ClpX proteases might be crucial for biofilm formation on glass by P482, when glycerol and citrate are present in the medium, for the cells to aggregate. Interestingly, the mutant in ClpA protease (KN1251) was capable of forming microcolonies in low-mass biofilm in the presence of citrate but not in glucose or glycerol. The *gacA* mutant, KN3318, analysed in this group of strains was able to form clustered biofilm with cells aggregating into microcolonies (Fig. 6A), however, in glucose it was more efficient than the wild-type P482 and in citrate its biofilm forming ability was decreased to about 50% of the wild-type.

**Figure 6.**
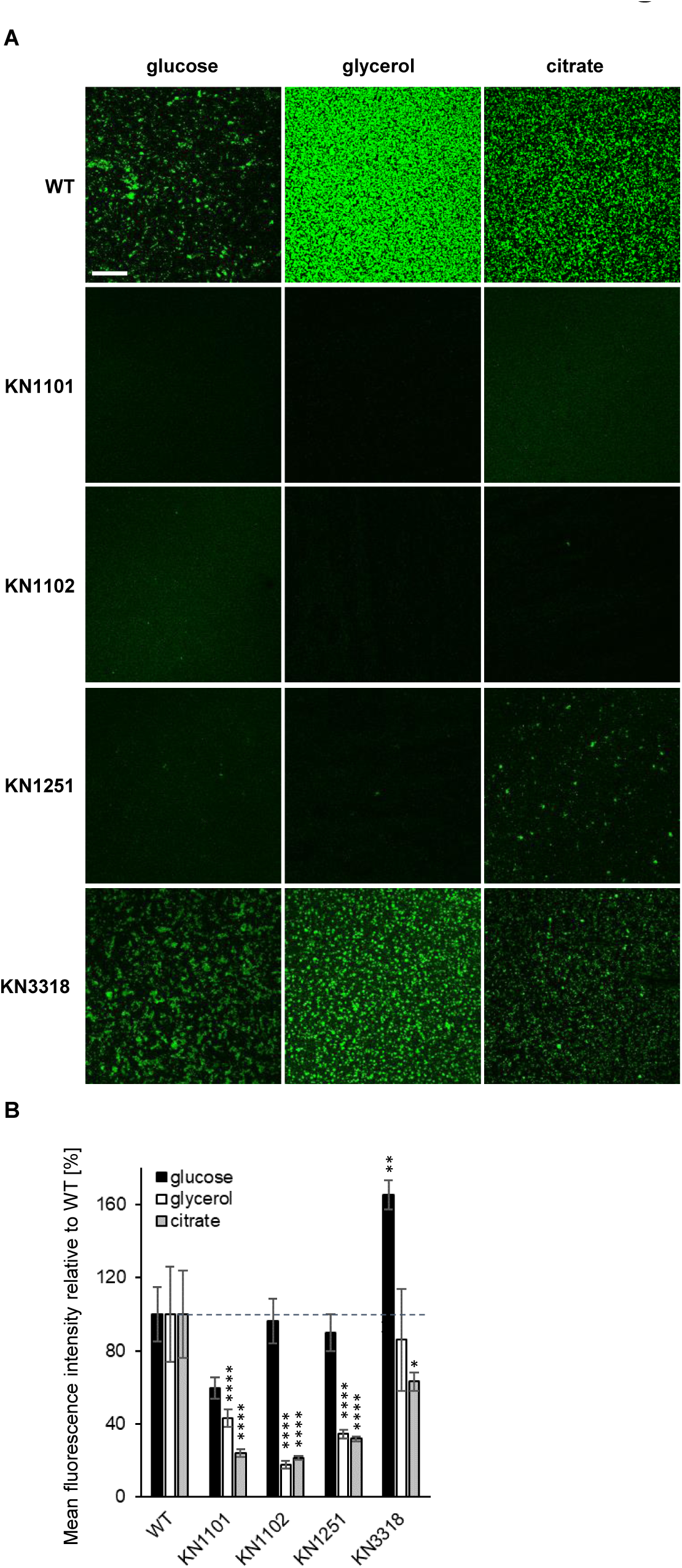
Biofilm formation on glass by GFP-tagged P482 mutants in proteases-encoding and *gacA* (KN3318) regulatory genes. (A) Representative CLSM images of biofilm formed by GFP-tagged P482 strains on the glass bottom of 24-well plates in M9 minimal medium supplemented with 0.4% glucose, 0.4% glycerol or 20 mM citrate. Bar = 200 µm. (B) Quantification of mean fluorescence intensity for the biofilm formed by the GFP P482 mutants in the M9 medium supplemented with the individual carbon sources. Statistical significance for mutants vs. WT was determined with ANOVA and Dunnet’s multiple comparisons test for correction (*P < 0.05; **P < 0.01; ***P < 0.001; ****P < 0.0001); all P values for the ANOVA analyses can be found in Supplementary Table S3.

In the group of mutants with knock-outs in matrix synthesis-related genes all occurred to be affected in biomass production in glycerol and citrate (Fig. 7A and B) in relation to the wild-type P482. The cells did attach to the surface of glass but managed to form only thin layer with very few, as in case of KN4700 and KN0543 mutants, or no, as in the remaining mutants, microcolonies scattered in the biofilm’s structure. Presence of glucose again did have an impact on the decrease of the strains’ ability to form more complex, three-dimensional biofilms, but allowed for accumulation of biomass, as reflected in the intensity of fluorescence signal recorded for the analysed z-stacks (Fig. 7B). This allows to assume that the potential of *P. donghuensis* P482 to form biofilm on the surface of glass greatly depends on its ability to synthesize the polysaccharide components of the biofilm matrix, particularly in medium where glycerol and citrate are the sole carbon sources.

**Figure 7.**
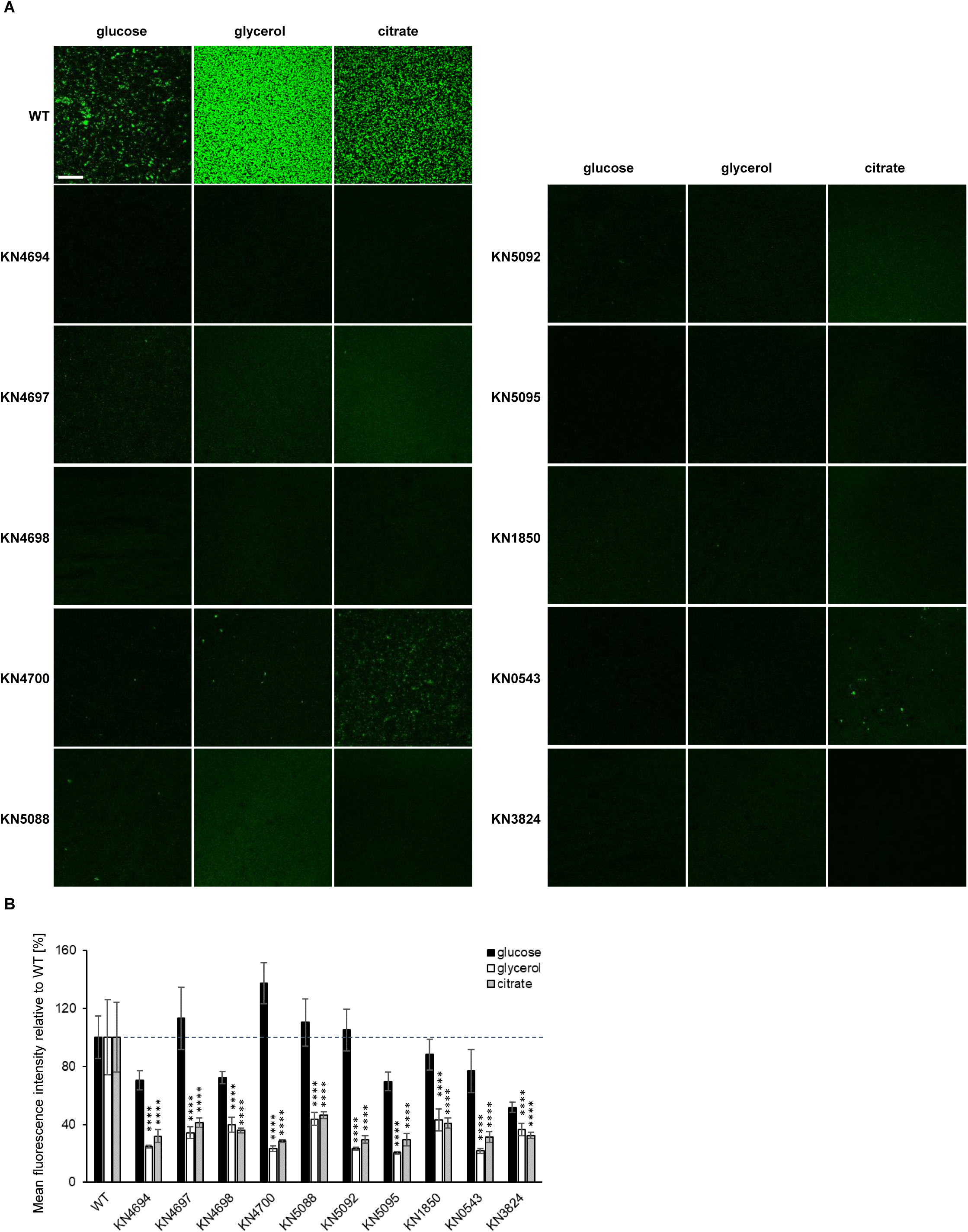
Biofilm formation on glass by GFP-tagged P482 mutants in matrix synthesis-related genes. (A) Representative CLSM images of biofilm formed by GFP-tagged P482 strains on the glass bottom of 24-well plates in M9 minimal medium supplemented with 0.4% glucose, 0.4% glycerol or 20 mM citrate. Bar = 200 µm. (B) Quantification of mean fluorescence intensity for the biofilm formed by the GFP P482 mutants in the M9 medium supplemented with the individual carbon sources. The bars represent means with SEM. Statistical significance for mutants vs. WT was determined with ANOVA and Dunnet’s multiple comparisons test for correction (*P < 0.05; **P < 0.01; ***P < 0.001; ****P < 0.0001); all P values for the ANOVA analyses can be found in Supplementary Table S3.

### The mutations in P482 genes do not affect colonization of plant roots of tomato or maize

Since during biofilm formation by the *P. donghuensis* P482 strains on the abiotic surfaces, polystyrene and glass, distinct phenotypes were observed, regulated by the available carbon sources and type of surface, it was only natural to ask whether the mutations could affect biofilm formation on biotic surface, that is, would they influence colonization of plant tissues? Two plant models were chosen for the analyses of plant tissues colonization as a form of biofilm formation on biotic surface by the P482, that is tomato and maize, as the representatives of the dicots and monocots, respectively. P482 is a tomato rhizosphere isolate and it has been shown previously to efficiently colonize other plant species, including maize (Krzyzanowska *et al*., 2012; Krzyżanowska *et al*., 2023).

The wild-type P482 strain colonized both types of plant rhizosphere efficiently, forming relatively large microcolonies scattered on the surface of tomato roots, and more abundant minute colonies on the surface of maize roots (Fig. 8, top left panel). The difference in the colonization pattern might had been due to the different culture conditions and dispersion of cells in the water environment in maize culture. Nevertheless, the P482 wild-type proved to efficiently colonize root tissue and persist on it for longer periods of time (one to four weeks, depending on the plant) in both experimental settings. The mutants with knock-outs in motility-associated genes demonstrated similar ability to attach to roots of both tomato and maize and form clusters of cells, as it was observed in case of the wild-type strain (Fig. 8), showing no defects in capacity to form biofilm on abiotic surface. Two of the mutants, KN0887 and KN0894 seemed to form numerous, more densely packed microcolonies on tomato roots, in the tested conditions, when the strains were used as inoculants individually.

**Figure 8.**
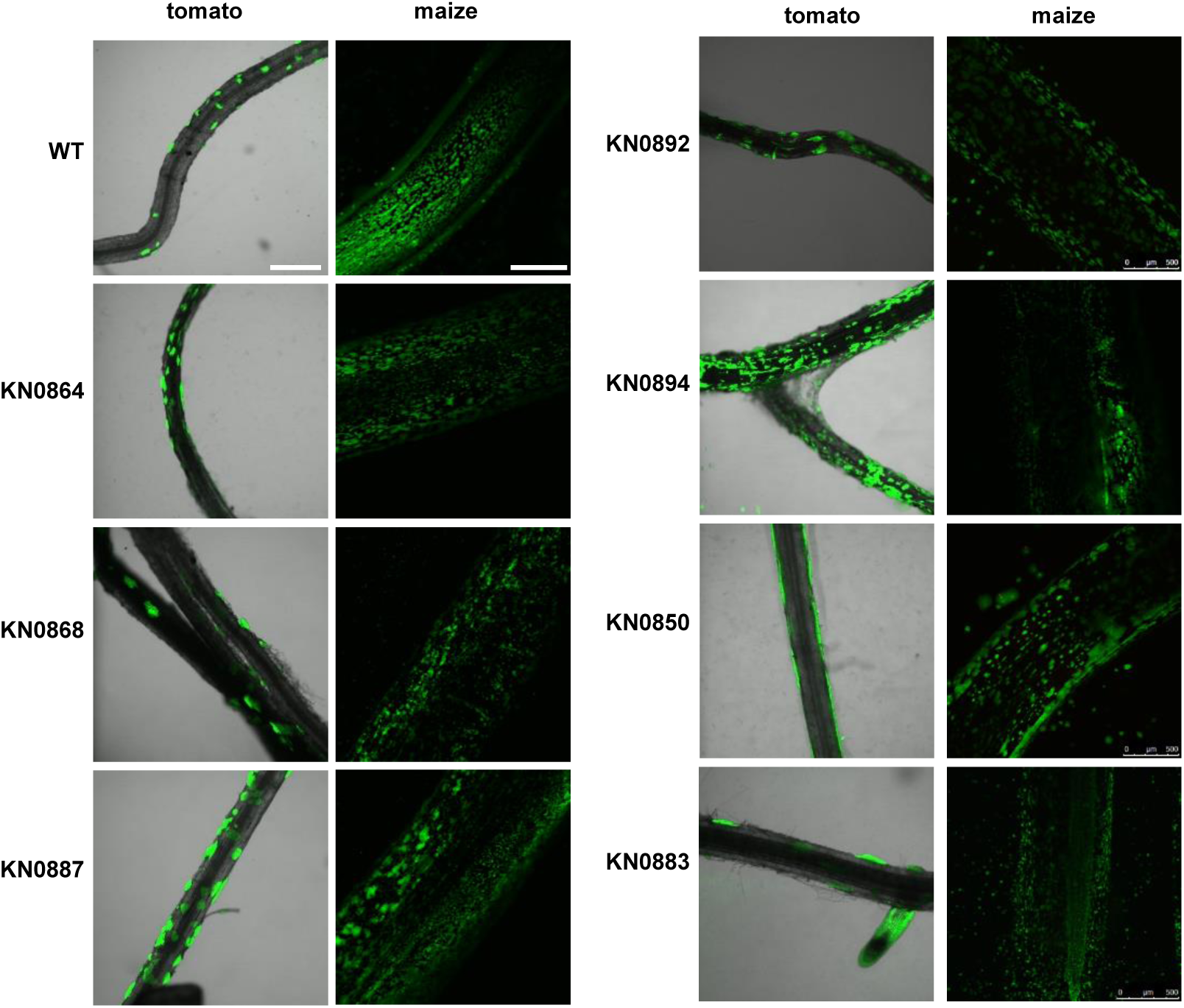
Colonization of plant rhizosphere by GFP-tagged P482 mutants in motility and attachment-related genes. Panels on the left show roots of tomato (merged bright-field and GFP channels), colonized with the P482 strains, visualized 4 weeks post inoculation; panels on the right show P482-colonized maize roots (GFP channel), visualized 1 week post inoculation (see Materials and Methods for details). Bars for tomato and maize = 500 µm.

Mutants in genes encoding proteases, KN1101, KN1102 and KN1251 (*clpP, clpX, clpA*, respectively), exhibited similar patterns and efficiency of colonization of the root tissue of tomato and maize as the wild-type strain (Fig. 9). KN1101 seemed to form slightly more microcolonies on the tomato roots than the wild-type did. The *gacA* mutant (KN3318) did not show any significant differences in tomato or maize rhizosphere colonization when compared to wild-type P482. This suggests that the mutations in proteases’ or the *gacA* regulatory genes are not affecting the biofilm formation by P482 on biotic surfaces. In the group of mutants with knock-outs in genes associated with synthesis of biofilm matrix components and adhesion three mutants, KN4700, KN5092 and KN5095, in genes encoding alginate-O-acetyltransferase, O-antigen ligase family protein and polysaccharide biosynthesis/export family protein, respectively, showed potential to be hypercolonizers of plant tissue, since the microcolonies were packed more densely on the surface of tomato roots (Fig. 10). The remaining mutants in this group behaved similarly to the wild-type strain, they formed scattered microcolonies on the tomato root tissue and multiple small colonies on maize roots. To sum up, the mutations introduced in the P482 genome, associated with different functions in the process of biofilm formation did not result in decrease of plant root colonization abilities. The strains demonstrated different phenotype when it comes to forming biofilm on biotic surface than what was observed on the abiotic surfaces, particularly glass.

**Figure 9.**
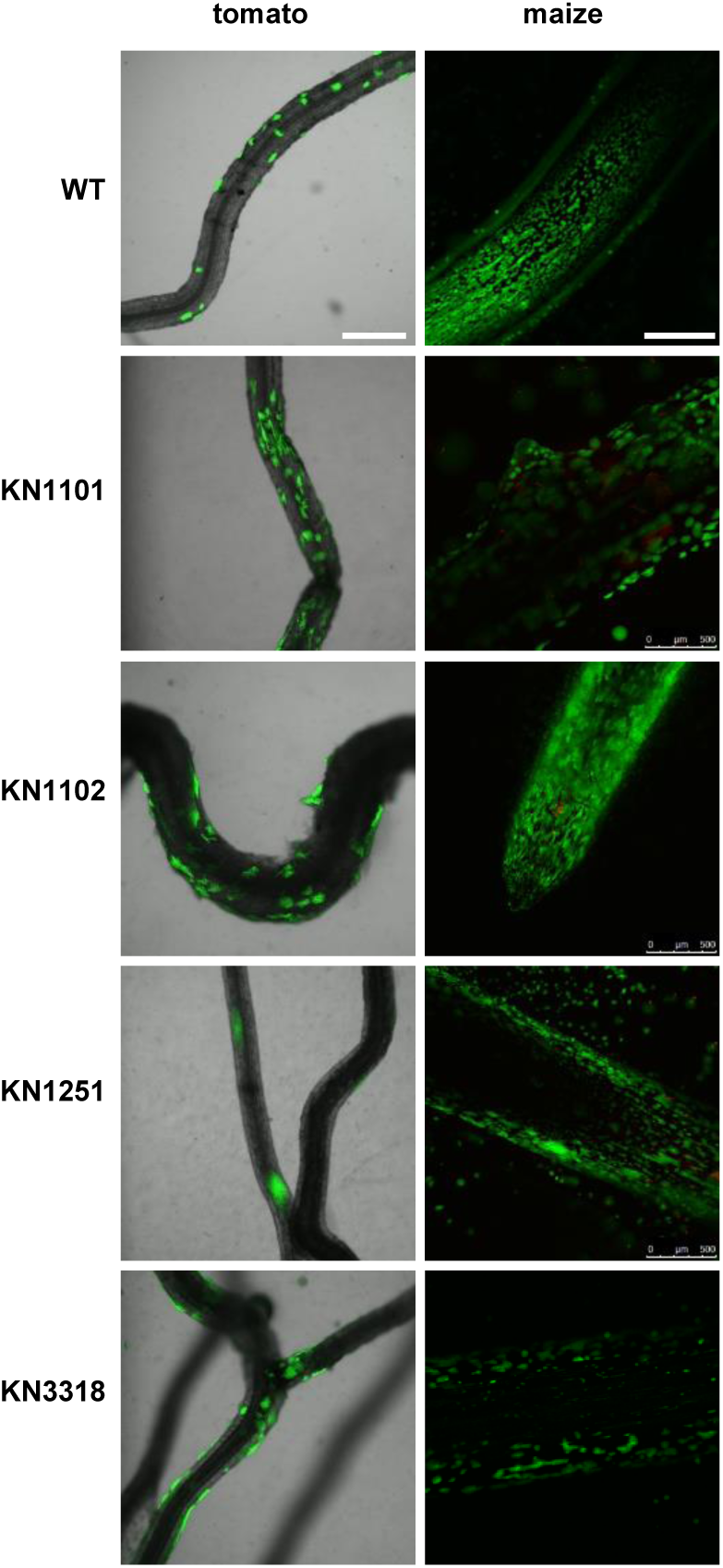
Colonization of plant rhizosphere by GFP-tagged P482 mutants in proteases-encoding and *gacA* (KN3318) regulatory genes. Panels on the left show roots of tomato (merged bright-field and GFP channels), colonized with the P482 strains, visualized 4 weeks post inoculation; panels on the right show P482-colonized maize roots (GFP channel), visualized 1 week post inoculation (see Materials and Methods for details). Bars for tomato and maize = 500 µm.

**Figure 10.**
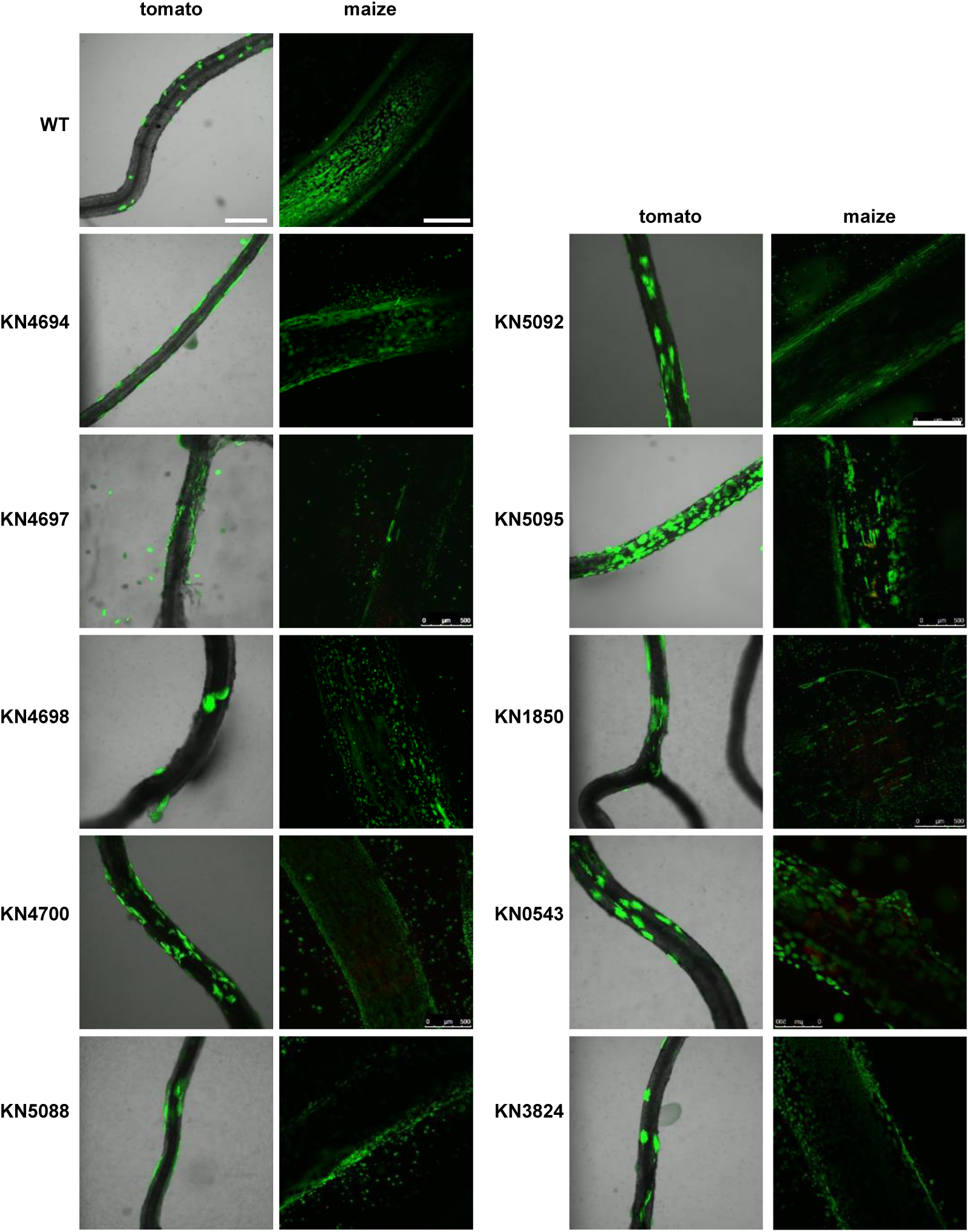
Colonization of plant rhizosphere by GFP-tagged P482 mutants in matrix synthesis-related genes. Panels on the left show roots of tomato (merged bright-field and GFP channels), colonized with the P482 strains, visualized 4 weeks aft post er inoculation; panels on the right show P482-colonized maize roots (GFP channel), visualized 1 week post inoculation (see Materials and Methods for details). Bars for tomato and maize = 500 µm.

## Discussion

The ability of plant-associated bacteria to form biofilm on the root tissues of their host is a crucial trait allowing them to persist in the rhizosphere environment and giving them advantage in the competition for nutrients (Ramey *et al*., 2004; Albareda *et al*., 2006; Danhorn and Fuqua, 2007; Espinosa-Urgel and Ramos-González, 2023). Many PGPR have been found to form some sort of biofilms on the roots of plants and the process has been studied widely for well-established species of Pseudomonads (Rudrappa, Splaine, *et al*., 2008; Barahona *et al*., 2010; Blanco-Romero *et al*., 2018; Zboralski and Filion, 2020; Costa-Gutierrez *et al*., 2022; Ajijah *et al*., 2023). Data concerning little known environmental isolates is scarce and the factors governing the mechanisms of biofilm formation and plant tissues colonization are not well understood. In this work the influence of carbon source and type of colonized surface, abiotic and biotic, on the biofilm-associated processes in the early stages of its formation were studied for the plant-beneficial *P. donghuensis* P482 wild-type strain and its knock-out mutants in biofilm-related genes.

The availability of particular sources of carbon, utilized by bacteria for growth and proliferation in the multicellular structures is one of the main environmental cues which play a role in the regulation of biofilm-associated processes. Many bacteria, including Pseudomonads, developed systems allowing them to select and assimilate various sources of carbon from the environment and manage their utilization through complex regulatory pathways: carbon catabolite repression, control and activation (CCR, CRC and CCA, respectively) systems (Deutscher, 2008; Görke and Stülke, 2008; Rojo, 2010). These contribute to optimization of bacterial metabolism in different circumstances, when nutrients are limited or in case of access to only one source of carbon, when there’s a need to utilize it in a balanced way to allow for most efficient growth (Bruckner and Titgemeyer, 2002). Strategies for carbon sources management in bacteria need to be therefore considered also in case of social behaviours, including biofilm formation (Kreft, 2004; Boyle *et al*., 2017). For *P. aeruginosa* it has been demonstrated that *crc* mutant was impaired in type IV pili synthesis, twitching motility and as a consequence also biofilm formation (O’toole *et al*., 2000).

For the wild-type strain of *P. donghuensis* P482 differences in the utilization of glucose, glycerol and citrate, used as sole carbon sources in the culture media, were reflected in the general production of biomass in the biofilm, as observed in crystal violet assay, and in the biofilm structure, visible in the CLSM images obtained for the GFP-tagged strain. Glycerol supplementation resulted in the most efficient biomass accumulation by the wild-type P482, despite the fact that this carbon source causes an extended lag phase during growth of the bacterium. What is more, in the presence of glycerol the strain formed the most developed, structured biofilm, consisting of three-dimensional clusters of microcolonies. It was reported for *P. aeruginosa* strains that glycerol metabolism promotes biofilm formation in PAO1, in comparison to when bacteria were grown on glucose (Scoffield and Silo-Suh, 2016). Mutation in *glpR*, which encodes transcriptional regulator managing the glycerol metabolism in *P. aeruginosa* resulted in increase of biofilm formation. In our work on antimicrobial properties of the P482 it was shown that glycerol impacts the expression of genes involved in synthesis of secondary metabolites in distinct way when compared to glucose (Matuszewska *et al*., 2021). Possible engagement of glycerol and other carbon sources in regulation of transcription of biofilm-associated genes in P482 is under investigation.

Different carbon sources available to *P. aeruginosa* during transition to sessile lifestyle resulted in diverse pattern of the biofilm – glucose promoted formation of more structured, three dimensional biofilm (Davies *et al*., 1998; Nivens *et al*., 2001) but the presence of citrate caused it to remain mostly flat (Heydorn *et al*., 2002; Klausen *et al*., 2003). The P482 biofilm structure did not reflect this observations; in glucose the biofilm formed by the wild-type P482 was the least developed of all three analysed carbon sources, and presence of citrate resulted in an intermediate type of biofilm. Glucose is not the preferred carbon source for *Pseudomonas* spp. and is metabolized fast when present individually in culture medium (Rojo, 2010). It does not cause lag phase during growth of P482 but influences wild-type biofilm formation negatively when compared to glycerol. It is plausible that different utilization of carbon sources in the P482 sessile and planktonic cells and a switch in metabolism control take place. It has been demonstrated for *Clostridium thermocellum* that sessile cells exhibit increased expression of genes involved in *i.a.* catabolism of carbohydrates by glycolysis and pyruvate fermentation and functions critical for cell division, and the planktonic cells have higher gene expression connected to e.g. motility and chemotaxis (Dumitrache *et al*., 2017). What needs to be taken into account is that the P482 biofilm analyses were performed at an early stage of formation and that its structure might evolve depending on the maturity of the biofilm as it was shown for example in *P. putida* strain grown in citrate (Tolker-Nielsen *et al*., 2000).

The P482 mutants demonstrated range of phenotypes depending on the carbon source supplemented during biofilm growth but also connected to the type of surface they colonized. For several mutants with knock-outs in motility or chemotaxis-associated genes glucose was utilized more efficiently than in the wild-type strain when it comes to biomass production when the cells attached to polystyrene or glass. Whereas for glycerol or citrate the strains produced biofilm similarly to the wild-type on polystyrene, but on glass almost all mutants showed significant decrease in ability to form biofilm. Active flagella did not seem to be required during attachment to polystyrene or glass when glucose was present in the minimal medium – the mutants accumulated biomass on both substrata equally well or with increased efficiency when compared to the wild-type strain. They were, however, unable to form microcolonies and larger structures as it was observed for the cells’ aggregation on glass. Glucose may act as an inhibitor of flagellar motility, as it was shown for *Vibrio vulnificus*, through dephosphorylation of EIIA^Glc^ (component of phosphoenolpyruvate: sugar phosphotransferase system, PTS) and sequestration of the FapA (flagellar assembly protein A) from the flagellated pole (Park *et al*., 2016). This was suggested as a mechanism allowing the bacteria to adapt to glucose-rich environments. What is more, the attachment in P482 might possibly be mediated by other structures than flagella but the accumulation of cells into clusters might depend on flagella-based scaffolding of the biofilm (Ozer *et al*., 2021). For various strains of *P. aeruginosa* it has been shown that the cells’ contact with surface may depend also on the motility facilitated by the activity of type IV pili (O’Toole and Kolter, 1998; Conrad, 2012) or that neither of the organelles are required for the cells’ attachment to abiotic surface as it was shown for PAO1 (Klausen *et al*., 2003; Burrows, 2012). The P482 genome contains numerous genes annotated as encoding type IV pilin proteins (e.g. *flp*), pilus assembly proteins (e.g. *pilX*) or type IV pili chemotaxis transducers, therefore it cannot be excluded that here they might act as appendages taking part in the early stages of biofilm formation and compensating for the absence or lack of activity of flagella.

The impact of nutrients and supplementation of media with different carbon sources was analysed for *P. fluorescens*, where glucose, citrate or glutamate were shown to stimulate biofilm formation through partially intersecting pathways (O’Toole and Kolter, 1998). The effect of carbon source was the more interesting since some of surface attachment deficient mutants (i.e. non-motile strains or *clpP* mutant) overcame the impact of genetic disturbance observed in glucose when supplemented with iron or grown in minimal medium containing citrate or glutamate as the sole carbon source. In P482 a similar mechanisms of compensation could take place, since some mutants, biofilm-deficient in the presence of glycerol or citrate (relatively to wild-type strain), were able to exhibit surface attachment and biomass production with glucose in the medium.

Formation of biofilm is regulated by different molecular mechanisms, for *Pseudomonas* and *Burkholderia* species they were specified as driven by cyclic diguanosine-5′-monophosphate (c-di-GMP), small RNAs (sRNA) or quorum sensing (QS) molecules (Fazli *et al*., 2014). The sRNAs system involves action of GacA/GacS/Rsm proteins, which among other modulate expression of motility, polysaccharide synthesis-related genes, or those with function in lipopeptide synthesis (Haas and Keel, 2003; Raaijmakers *et al*., 2010; Duque *et al*., 2013). However, the effects of mutations in the genes encoding the components of this regulatory system seem to depend on a bacterial strain. Increased expression of flagellin and enhanced motility have been observed in *gacA* mutants of root-colonizing *P. fluorescens* F113 (Sánchez-Contreras *et al*., 2002). In *P. aeruginosa* PAO1 *gacA* mutant increased swarming and biofilm formation were observed (Kay *et al*., 2006). The P482 *gacA* mutant exhibited non-motile phenotype in swarming tests and showed decreased synthesis of matrix polysaccharides as seen in the colony morphology test, but its ability to form biofilm on abiotic or biotic surfaces was mostly unaffected and even surpassed the wild-type strain during attachment to polystyrene in the presence of all tested carbon sources, indicating that biofilm formation is controlled negatively by this system in *P. donghuensis*. On glass the *gacA* mutant cells aggregated into microcolonies more efficiently than the wild-type when glucose was sole carbon source but less so in presence of citrate in the medium. We observed that *gacA* expression in wild-type P482 was affected negatively in medium supplemented with glycerol, in contrast to that with glucose (Matuszewska *et al*., 2021).

A possible pathway of biofilm control was proposed for *P. fluorescens* and includes the activity of Clp proteases (O’Toole and Kolter, 1998), where a motile *clpP* mutant showed defects in biofilm formation on both hydrophilic and hydrophobic surfaces. For *P. aeruginosa* PAO1 the Lon protease was shown to be essential for biofilm formation (Marr *et al*., 2007), through a possible involvement in protein degradation in order to increase amino acid pool available during starvation, what occurs in case of *Escherichia coli* or *Salmonella typhimurium* (Miller, 1996). ClpP, ClpX and ClpA in P482, being subunits of an ATP-dependent Clp protease, also appear to be engaged in biofilm formation processes, as the activity of mutants in the genes encoding them varied, depending on the surface and carbon source tested. Glucose and citrate seemed to compensate the mutation in *clpX* gene on polystyrene, but on glass the citrate and glycerol had negative impact on the formation of structured biofilm. The role of Clp proteases in biofilm control may be attributed to their general activity, that is degradation of misfolded proteins, transcription factors and proteins involved in cellular signalling. In *Bacillus subtilis* Clp proteases down-regulate the central metabolic pathways when cells enter state of glucose starvation (Gerth *et al*., 2008).

Biofilm matrix polysaccharides synthesized by bacteria enable their survival and persistence in various niches, through protection from external factors. The polysaccharide biosynthesis involves multiple energy-requiring processes and regulation (Flemming and Wingender, 2010; Flemming *et al*., 2023), when various carbohydrates (mannose, glucose, rhamnose, mannuronic acid, etc.) are incorporated into the polymeric chains (Ma *et al*., 2006; Vassoler Serrato, 2022). *P. donghuensis* P482 genome encodes numerous enzymes involved in the synthesis of alginate, undefined exopolysaccharides, LPS and O-antigen, and also enzymes catalysing the cleavage reactions of the polymers. In our experimental setup knock-out mutations in the genes encoding enzymes involved in biosynthesis of matrix components in P482 had little impact on biofilm formation on polystyrene, regardless of carbon source utilized by the cells. Only in case of two genes, encoding enzymes engaged in alginate synthesis (alginate production protein AlgE and membrane bound alginate O-acetyltransferase) increased biofilm production was observed in the presence of glucose, in relation to the wild type strain. Change of substratum to glass, resulted in lower biofilm formation for all strains lacking the inactivated enzymes associated with matrix compounds biosynthesis in the presence of glycerol and citrate, when compared to wild-type, but the inability to develop biofilm structures was observed for all three carbon sources. It has been shown for *P. aeruginosa* that mutations in genes involved in the synthesis of exopolysaccharide alginate or LPS affect also the twitching motility (Whitchurch *et al*., 1996; Abeyrathne *et al*., 2005), impacting bacterial attachment to surfaces. Notwithstanding, the inactivation of enzymes regulating biosynthesis of polysaccharides mostly hinders efficient production of biofilm matrix, what prevents cells from attaching to surfaces and aggregating into larger structures. It has been shown for *P. aeruginosa* that glucose efficiently promotes biofilm formation by upregulation of expression of the *pslA* gene, related to biosynthesis of extracellular polysaccharide (She *et al*., 2019). Mutation in *glpR* regulator, which represses expression of genes involved in glycerol metabolism also enhances biofilm formation through upregulation of Pel polysaccharide (Scoffield and Silo-Suh, 2016). However, in *P. aeruginosa* evolution experiments on *psl* and *pel* mutants it was observed that inactivation of one polysaccharide synthesis gene leads to defects in initial biofilm formation but can promote expression of compensatory matrix polysaccharides with longer time of biofilm growth (Colvin *et al*., 2012). Extended biofilm growth experiments and gene expression analyses could therefore shed more light on the regulation of polysaccharide synthesis in biofilm matrix in P482. Also, since the exact composition of matrix polysaccharides in P482 has not been studied up to date, it would be worth an attempt to isolate and characterize them for the wild-type and the mutant strains.

The results obtained for the two types of abiotic surfaces used in the experiments in this work allow to think that also the character of the surface had an impact on the interactions between bacterial cells and the colonized environment. Physicochemical features of the artificial substrata such as surface charge, hydrophobicity or surface roughness do affect bacterial attachment (Palmer *et al*., 2007). The P482 mutants, irrespective of the type of introduced mutation, generally attached more efficiently to hydrophobic polystyrene than to glass, which is more hydrophilic. Bacterial cells, due to the composition of their cell wall molecules, e.g. proteins and lipids, and the character of polymeric substances surrounding them in biofilm matrix tend to be more hydrophobic in nature (Davies *et al*., 1998; Zheng *et al*., 2021) and therefore more prone to attach to hydrophobic surfaces. For *Escherichia coli* it was observed that moderate substrate hydrophobicity provides the highest adhesion of cells (Yuan *et al*., 2017). Hydrophilic surfaces (e.g. borosilicate glass) affected negatively the ability of *P. aeruginosa* or *Staphylococcus epidermidis* to form biofilms (De-la-Pinta *et al*., 2019), but also superhydrophobic polymers in case of *S. aureus, P. aeruginosa* or *E. coli* biofilms significantly reduced the cellular attachment and delayed it in time (Loo *et al*., 2012; Ozkan *et al*., 2020; Montgomerie and Popat, 2021; Zheng *et al*., 2021). This trait might be of importance in case of bacterial attachment to plant surfaces, including root tissue. The hydrophobicity of the maize root mucilage has been suggested to also contribute to the efficient attachment of bacteria to the plant (Holz *et al*., 2018). Attachment of *P. fluorescens* F113 strains to roots of alfalfa was possible, among other factors, due to a plant-derived mucigel which holds bacteria forming microcolonies on the surface of the plant tissue (Barahona *et al*., 2010). Similar results were obtained for *P. fluorescens* CHA0, *P. putida*, *Xanthomonas oryzae* or *Acidovorax facilis* colonizing tomato tissue (Chin-A-Woeng *et al*., 1997).

The microbial ability to form biofilm on abiotic surfaces and colonization of plant tissues do not, however, simply go hand in hand, and depend largely on the bacterial species. The complexity of bacterial adaptations to sessile lifestyle is reflected in the variety of mechanisms which lead to formation of biofilms on abiotic and biotic surfaces (Danhorn and Fuqua, 2007; Espinosa-Urgel and Ramos-González, 2023). In *P. putida* for example the flagella and systems of synthesis and transport of large adhesin LapA play a role in attachment to both artificial surfaces and plant seeds, but impairment of synthesis of LPS was crucial only for seed and root colonization but not attachment to plastic (Yousef-Coronado *et al*., 2008). In *P. fluorescens* a surface attachment defective mutant showed reduced biofilm formation on abiotic surface but did not exhibit impairment in alfalfa root colonization (Barahona *et al*., 2010). The case that all strains of *P. donghuensis* P482, wild-type and mutants, did settle on the root tissues of both tomato and maize efficiently but showed different biofilm formation abilities depending on the abiotic surface in combination with particular mutations indicates composite and alternative mechanisms governing these processes in this species. Also, the strains were analysed individually in the colonization assays and it would be interesting to verify their colonization abilities in competition tests, where the wild-type P482 would also be present. It was shown for *P. fluorescens* F113 that non-motile mutants which colonized alfa-alfa roots when tested individually, were poor competitors when inoculated together with the wild-type strain (Capdevila *et al*., 2004).

Moreover, the ability of all P482 mutants to colonize plant roots can be attributed to the fact that not only bacterial factors take part in the interactions between the cells and the colonized biotic surface provided by the host. For various bacterial species it has been demonstrated that the root exudates and substances secreted by the plant cells also play role during establishment of microbiome on the host tissues (Danhorn and Fuqua, 2007; Rudrappa, Biedrzycki, *et al*., 2008; Rudrappa, Splaine, *et al*., 2008). In a recent work by Krzyżanowska and colleagues (Krzyżanowska *et al*., 2023) it has been demonstrated that P482 exhibits different patterns of gene expression in response to root exudates of the two plant hosts, tomato and maize. Induction of genes associated in P482 with motility, e.g. *fimV* encoding protein involved in the assembly of type IV pili or *fliS* encoding one of flagellar proteins, took place under the influence of maize exudates, in contrast to reduction of expression of genes associated with chemotaxis by tomato exudates. The plant root metabolites often serve as a trigger signal for the bacteria to colonize plant tissues and form microcolonies at sites where the exudates are released from cells to the soil (Lugtenberg *et al*., 1999; Rudrappa, Splaine, *et al*., 2008) but can also act as biofilm formation inhibitors in relation to bacterial plant pathogens (Walker *et al*., 2004). An important trait in tomato root colonization by *P. fluorescens* WCS365 was the flagella-driven chemotaxis towards exudate components (De Weert *et al*., 2002).

The effect of specific environmental cues on the individual processes involved in bacterial transition to sessile lifestyle still requires multi-level investigations but the results obtained in this work add to the notion that the biofilm formation depends on intricate sequence of events leading to construction of multicellular bacterial community in the most favourable environment. To conclude, the complexity of mechanisms involved in plant host colonization and biofilm formation by less-known strains such as *P. donghuensis* P482 still requires detailed research, since many factors, both cellular and environmental, play significant roles in the processes. Elucidating them is of particular importance when plant growth promoting and biocontrol bacteria are concerned. Their eventual use in field applications must take into account their long-term survival in the target environment what is more certain in case of biofilm-encased than planktonic cells. Future research could focus on investigating possible compensation of mutation-affected pathways by alternative cellular mechanisms, leading to efficient establishment of bacterial on plant tissues.

## Acknowledgements

The authors would like to thank Prof. Paul Williams (Centre for Biological Sciences, University of Nottingham, United Kingdom) for his valuable support and advising during the course of the project. We thank Mikołaj Pęczak for technical support with motility in medium S assay. We also highly appreciate the input of the anonymous Reviewers.

## Author contributions

MR conceived the study, acquired funding, performed the experimental work, analyzed all data, prepared all figures and tables, and drafted the manuscript. TM constructed KN0868 and KN4698 mutant strains. SJ supervised and coordinated the project, and revised the manuscript.

## Competing interests

The authors declare no competing interests.

**Correspondence** and requests for materials should be addressed to S.J.

