## Supplementary Figures for "Carbon source and surface type influence the early-stage biofilm formation by rhizosphere bacterium *Pseudomonas donghuensis* P482"

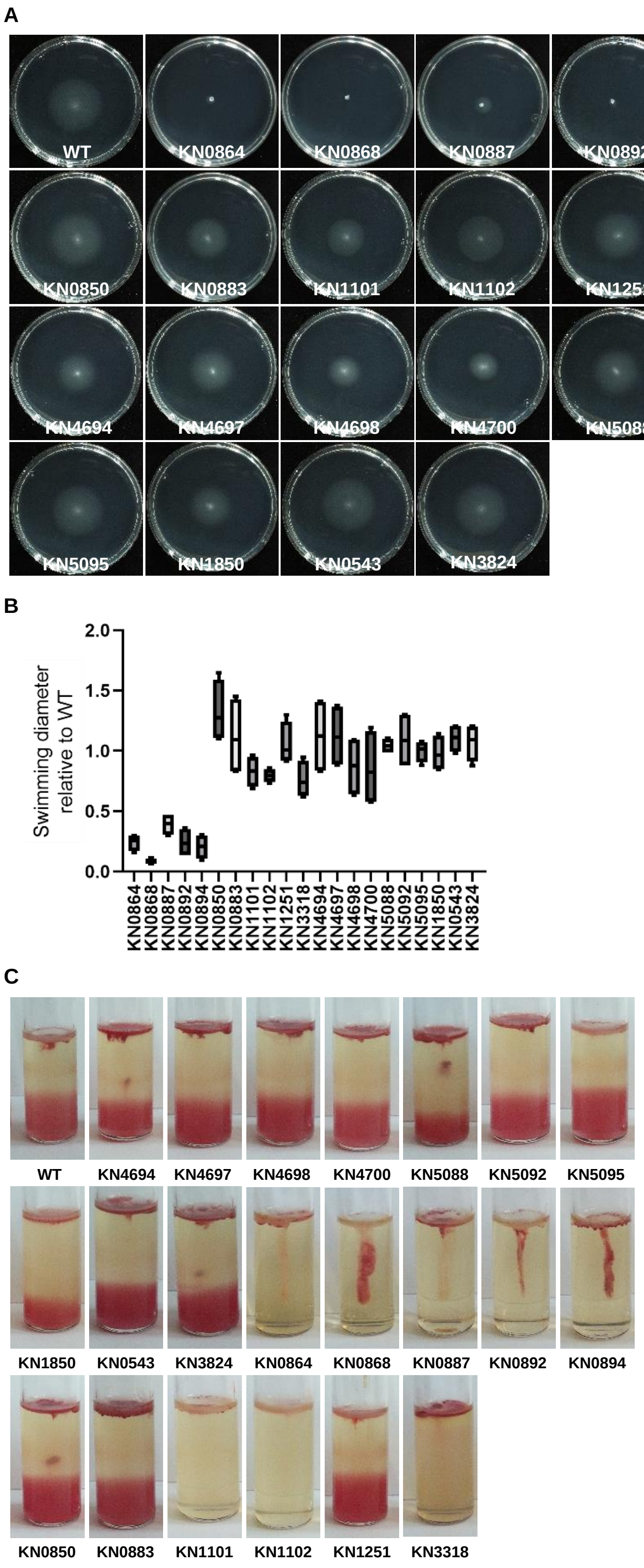


**Supplementary Figure S1. Swimming motility data for all analysed P482 strains.** (A) Representative images of all analysed P482 mutants versus the wild-type strain. (B) Quantification of swimming diameter of all analysed mutants relative to the wild-type strain (taken as 1), n=4. (C) Results of motility assay in motility S medium for all analysed P482 mutants.

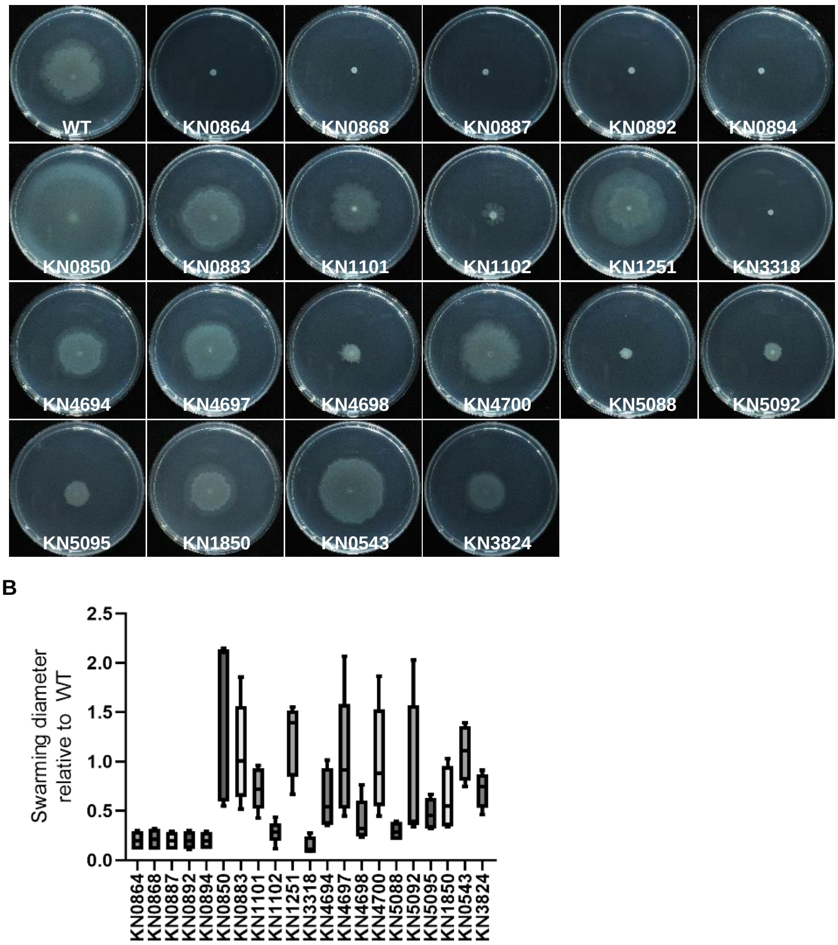


**Supplementary Figure S2. Swarming motility data for all analysed P482 strains.** (A) Representative images of all analysed P482 mutants versus the wild-type strain. (B) Quantification of swarming diameter of all analysed mutants relative to the wild-type strain, n=4 or 5.

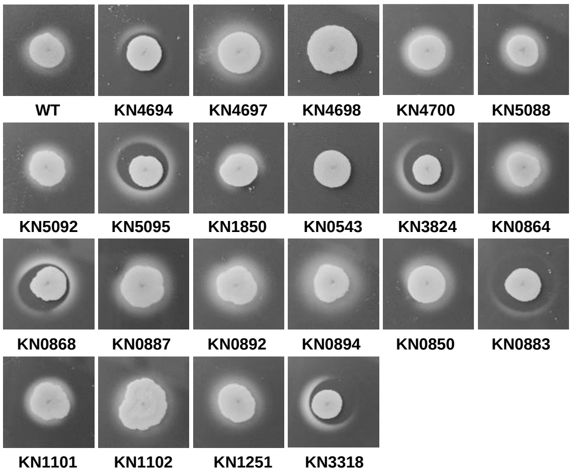


**Supplementary Figure S3. Twitching motility data for all analysed P482 strains.** Images of all P482 strain colonies with twitching halos, shown in black and white for better contrast. The mutants exhibiting defect in the twitching motility were KN4698 and KN0543.

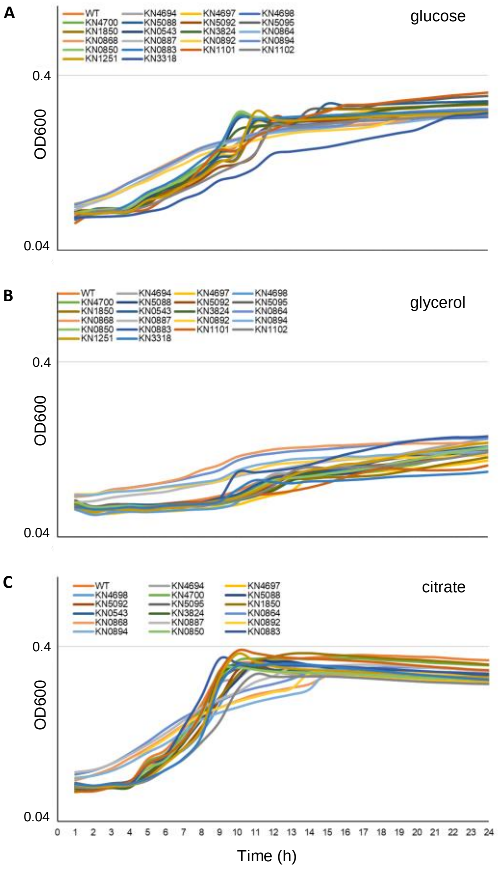


**Supplementary Figure S4. Growth curves for *P. donghuensis* P482 wild type strain and its mutants in minimal medium supplemented with different carbon sources.** The strains were cultured in M9 medium supplemented with (A) 0.4% glucose, (B) 0.4% glycerol or (C) 20 mM citrate, in 96-well plate in 3 technical replicates each, for 24 h at 28°C, with shaking. The 600 nm absorbance measurements ($OD_{600}$) were done every 20 minutes. Mean value for each readout was used to produce growth curves of each strain under given conditions.
