## Supplementary Tables for "Carbon source and surface type influence the early-stage biofilm formation by rhizosphere bacterium *Pseudomonas donghuensis* P482"

**Supplementary Table S1.** PCR primers designed and used in this study.

| **Locus** | **Name** | **Sequence** | **Introduced cloning site** | **Amplicon length (bp)** |
|---|---|---|---|---|
| BV82_0864 | XbaI_P482_0864_F | GCGCGC**TCTAGA**CGCAGCCAGACTACGATGAC | XbaI | 473 |
| | ApaI_P482_0864_R | GCGCGC**GGGCCC**GGCGGTCTTGAACTTGATGAC | ApaI | |
| BV82_0868 | XbaI_P482_0868_F | TTCGC**TCTAGA**CCACACCCTGGACAAGTTCT | XbaI | 400 |
| | Xho_P482_0868_R | TTAAA**CTCGAG**GAGCATGGTCAGGAACAGGT | XhoI | |
| BV82_0887 | XbaI_P482_0887_F | GCGCGC**TCTAGA**CATGTTCAACCTGCTGCGTC | XbaI | 460 |
| | Xho_P482_0887_R | GCGCGC**CTCGAG**TCATCGAGTACGGCATGGTCAC | XhoI | |
| BV82_0892 | XbaI_P482_0892_F | GCGCGC**TCTAGA**CGCATTCGCCTGTATTTCGCC | XbaI | 481 |
| | Kpn_P482_0892_R | GCGCGC**GGTACC**AAAGCAATGTTGACCACCAGCAG | KpnI | |
| BV82_0894 | XbaI_P482_0894_F | GCGCGC**TCTAGA**CAACATGACCTTTGCCGATGCC | XbaI | 451 |
| | Xho_P482_0894_R | GCGCGC**CTCGAG**GAATCAGCCGATAGCCGACC | XhoI | |
| BV82_0850 | XbaI_P482_0850_F | GCGCGC**TCTAGA**CGGCATCAACGTGTTCAAGG | XbaI | 463 |
| | Kpn_P482_0850_R | GCGCGC**GGTACC**AGTCATCGACCGTCAGCACC | KpnI | |
| BV82_0883 | XbaI_P482_0883_F | GCGCGC**TCTAGA**ATCACAATCACCTGCTCAACGA | XbaI | 542 |
| | Kpn_P482_0883_R | GCGCGC**GGTACC**ACAGCAAGAACAACCGCTCTC | KpnI | |
| BV82_1101 | XbaI_P482_1101_F | GCGCGC**TCTAGA**AGCGGGTGATCTTCCTGGTC | XbaI | 428 |
| | Kpn_P482_1101_R | GCGCGC**GGTACC**CAGTGTCACGCTTGATGGTCTC | KpnI | |
| BV82_1102 | XbaI_P482_1102_F | GCGCGC**TCTAGA**GTACAACCACTACAAGCGCCT | XbaI | 476 |
| | Kpn_P482_1102_R | GCGCGC**GGTACC**GCACCGCCACAGATGAACAG | KpnI | |
| BV82_1251 | XbaI_P482_1251_F | GCGCGC**TCTAGA**GCGGCATCTTCGAGAAAGACC | XbaI | 574 |
| | Xho_P482_1251_R | GCGCGC**CTCGAG**GAAACGCACCAGCTCAACCC | XhoI | |
| BV82_4694 | XbaI_P482_4694_F | GCGCGC**TCTAGA**CGGCTACCGACATCATCGAGAC | XbaI | 520 |
| | Kpn_P482_4694_R | GCGCGC**GGTACC**GTTGTAGGCGTCGCTGTTGG | KpnI | |
| BV82_4697 | XbaI_P482_4697_F | GCGCGC**TCTAGA**GATTCCACCGCGAGCATCAC | XbaI | 527 |
| | Kpn_P482_4697_R | GCGCGC**GGTACC**CGGCAGGGCATAGTTGTGGT | KpnI | |

| | | | | |
|---|---|---|---|---|
| BV82_4698 | XbaI_P482_4698_2_F | ATTAT**TCTAGA**ACGGTGCCAACTTCACCTAC | XbaI | 445 |
| | Xho_P482_4698_2_R | AATTC**CTCGAG**AGAATCGAGGCAACGAACAG | XhoI | |
| BV82_4700 | XbaI_P482_4700_F | GCGCGC**TCTAGA**CCGAAAGGCTCGACCTTCGT | XbaI | 392 |
| | Kpn_P482_4700_R | GCGCGC**GGTACC**GCTGACCTTGACCGGATTGATCTC | KpnI | |
| BV82_5088 | XbaI_P482_5088_F | GCGCGC**TCTAGA**ATCAGCCACTCGCCGATCAG | XbaI | 624 |
| | Kpn_P482_5088_R | GCGCGC**GGTACC**GCAACGAACTTGAAGTGACCACC | KpnI | |
| BV82_5092 | XbaI_P482_5092_F | GCGCGC**TCTAGA**CGTGTTTGCCATGTTCATCGCC | XbaI | 400 |
| | Kpn_P482_5092_R | GCGCGC**GGTACC**GGATCATCTTGCTGCCGACC | KpnI | |
| BV82_5095 | XbaI_P482_5095_F | GCGCGC**TCTAGA**ACTTGAGCATCACGTCCACGG | XbaI | 476 |
| | Xho_P482_5095_R | GCGCGC**CTCGAG**TCTTCCACATCAGTACCAGCGG | XhoI | |
| BV82_1850 | XbaI_P482_1850_F | GCGCGC**TCTAGA**CGACTTGCACCTGGACAACCT | XbaI | 492 |
| | Kpn_P482_1850_R | GCGCGC**GGTACC**GATGTTATAGCGTTTGAGCACCAG | KpnI | |
| BV82_0543 | XbaI_P482_0543_F | GCGCGC**TCTAGA**CGAAGCCAGGTTCTATGAAGAG | XbaI | 453 |
| | Xho_P482_0543_R | GCGCGC**CTCGAG**GAGAATCCAGTAGGTCAGCGG | XhoI | |
| BV82_3824 | XbaI_P482_3824_F | GCGCGC**TCTAGA**CATGTTGATGATGCCCTGGT | XbaI | 482 |
| | Kpn_P482_3824_R | GCGCGC**GGTACC**TTCACAGGTCAATTCGAGGAG | KpnI | |

The listed oligonucleotides were synthesized by Sigma-Aldrich (USA). Sequences recognized by respective restriction enzymes are given in bold. Annealing temperature for all primers was 65°C.

**Supplementary Table S2.** P values from ANOVA analyses for the listed tests for the *P. donghuensis* P482 mutants' biofilm phenotypes compared to the wild-type strain.

| Comparison | Swimming | Swarming | Polystyrene (glucose) | Polystyrene (glycerol) | Polystyrene (citrate) | Glass (glucose) | Glass (glycerol) | Glass (citrate) |
|---|---|---|---|---|---|---|---|---|
| WT vs. KN0864 | **<0.0001**[d] | **0,0356**[a] | 0,9955 | **<0.0001**[d] | 0,9991 | **<0.0001**[d] | **<0.0001**[d] | **<0.0001**[d] |
| WT vs. KN0868 | **<0.0001**[d] | **0,0392**[a] | 0,1385 | 0,6291 | 0,9168 | **<0.0001**[d] | **<0.0001**[d] | **<0.0001**[d] |
| WT vs. KN0887 | **0,0004**[c] | **0,0342**[a] | **0,0175**[a] | 0,3858 | 0,1005 | 0,5725 | **<0.0001**[d] | **<0.0001**[d] |
| WT vs. KN0892 | **<0.0001**[d] | **0,0345**[a] | **0,0004**[c] | 0,9654 | **0,0043**[b] | 0,9999 | **0,0002**[c] | 0,9955 |
| WT vs. KN0894 | **<0.0001**[d] | **0,0343**[a] | **<0.0001**[d] | 0,1888 | **0,0002**[c] | **0,017**[a] | 0,3811 | 0,9994 |
| WT vs. KN0850 | 0,1783 | 0,3357 | 0,0588 | 0,955 | 0,999 | **<0.0001**[d] | 0,9872 | **<0.0001**[d] |
| WT vs. KN0883 | 0,9943 | 0,9996 | 0,5557 | 0,2498 | >0,9999 | 0,9585 | **<0.0001**[d] | **<0.0001**[d] |
| WT vs. KN1101 | 0,9023 | 0,9695 | 0,9945 | 0,1953 | 0,9883 | 0,2403 | **<0.0001**[d] | **<0.0001**[d] |
| WT vs. KN1102 | 0,7487 | 0,0538 | **<0.0001**[d] | 0,9957 | **0,0004**[c] | 0,9998 | **<0.0001**[d] | **<0.0001**[d] |
| WT vs. KN1251 | 0,9994 | 0,9884 | 0,9998 | 0,0715 | 0,9997 | 0,9993 | **<0.0001**[d] | **<0.0001**[d] |
| WT vs. KN3318 | 0,5542 | **0,0113**[a] | **<0.0001**[d] | **<0.0001**[d] | **<0.0001**[d] | **0,0027**[b] | 0,9511 | **0,0111**[a] |
| WT vs. KN4694 | 0,9899 | 0,7851 | **0,0199**[a] | 0,9158 | 0,9995 | 0,7307 | **<0.0001**[d] | **<0.0001**[d] |
| WT vs. KN4697 | 0,9896 | 0,9999 | 0,1164 | **0,0263**[a] | >0,9999 | 0,9991 | **<0.0001**[d] | **<0.0001**[d] |
| WT vs. KN4698 | 0,9878 | 0,1696 | **<0.0001**[d] | 0,9993 | 0,9991 | 0,8166 | **<0.0001**[d] | **<0.0001**[d] |
| WT vs. KN4700 | 0,9733 | >0,9999 | 0,8542 | 0,9996 | 0,9995 | 0,3616 | **<0.0001**[d] | **<0.0001**[d] |
| WT vs. KN5088 | 0,9996 | 0,0893 | 0,5818 | **0,0049**[b] | 0,9997 | 0,9993 | **<0.0001**[d] | **<0.0001**[d] |

| | | | | | | | | |
|---|---|---|---|---|---|---|---|---|
| WT vs. KN5092 | 0,999 | 0,9991 | 0,1369 | 0,7277 | 0,832 | 0,9997 | **<0.0001**[d] | **<0.0001**[d] |
| WT vs. KN5095 | >0,9999 | 0,3047 | 0,9946 | 0,9949 | 0,9521 | 0,6909 | **<0.0001**[d] | **<0.0001**[d] |
| WT vs. KN1850 | 0,9998 | 0,7946 | 0,9248 | 0,1485 | 0,522 | 0,9992 | **<0.0001**[d] | **<0.0001**[d] |
| WT vs. KN0543 | 0,9956 | 0,9996 | 0,3433 | 0,9991 | 0,9994 | 0,9496 | **<0.0001**[d] | **<0.0001**[d] |
| WT vs. KN3824 | 0,9993 | 0,9751 | 0,117 | 0,9991 | 0,9993 | 0,0763 | **<0.0001**[d] | **<0.0001**[d] |

Statistically significant P values are marked in bold.

[a] – P value < 0.05, [b] - P value < 0.01, [c] – P value < 0.0005, [d] – P value < 0.0001

**Supplementary Table S3.** P values from ANOVA analyses comparing efficiency of biofilm formation on abiotic surfaces (polystyrene or glass) for each tested *P. donghuensis* P482 strain in the three tested carbon sources.

| | Biofilm on polystyrene | | | Biofilm on glass | | |
|---|---|---|---|---|---|---|
| *P. donghuensis* P482 strain | glucose vs. glycerol | glucose vs. citrate | glycerol vs. citrate | glucose vs. glycerol | glucose vs. citrate | glycerol vs. citrate |
| WT | **0.0002**[c] | 0.0562 | **<0.0001**[d] | **<0.0001**[d] | **0.0142**[a] | **0.0011**[b] |
| KN0864 | 0.057 | **0.0004**[c] | **<0.0001**[d] | 0.0879 | **0.0021**[b] | 0.228 |
| KN0868 | **<0.0001**[d] | **<0.0001**[d] | **<0.0001**[d] | 0.2253 | **0.0084**[b] | 0.2551 |
| KN0887 | **<0.0001**[d] | **<0.0001**[d] | **<0.0001**[d] | 0.065 | **0.0349**[a] | 0.9513 |
| KN0892 | **<0.0001**[d] | **<0.0001**[d] | **<0.0001**[d] | 0.2016 | **0.0205**[a] | 0.4871 |
| KN0894 | 0.5025 | **<0.0001**[d] | **<0.0001**[d] | 0.1446 | 0.85 | 0.3469 |
| KN0850 | **<0.0001**[d] | **0.0005**[c] | **<0.0001**[d] | 0.8393 | **<0.0001**[d] | **<0.0001**[d] |
| KN0883 | **<0.0001**[d] | **<0.0001**[d] | **<0.0001**[d] | **0.0478**[a] | 0.1169 | **0.0004**[c] |
| KN1101 | **<0.0001**[d] | 0.1776 | **<0.0001**[d] | **<0.0001**[d] | 0.2992 | **<0.0001**[d] |
| KN1102 | **0.0008**[c] | **0.0013**[b] | **<0.0001**[d] | **<0.0001**[d] | **<0.0001**[d] | 0.2158 |
| KN1251 | **<0.0001**[d] | **0.0352**[a] | **<0.0001**[d] | 0.1386 | **0.0018**[b] | **<0.0001**[d] |
| KN3318 | **<0.0001**[d] | >0.9999 | **<0.0001**[d] | 0.1521 | 0.6253 | **0.0236**[a] |
| KN4694 | **0.0001**[c] | **<0.0001**[d] | **<0.0001**[d] | 0.7781 | 0.2681 | 0.0813 |
| KN4697 | **<0.0001**[d] | **0.0264**[a] | **<0.0001**[d] | 0.8176 | 0.057 | 0.1785 |
| KN4698 | **<0.0001**[d] | **<0.0001**[d] | **<0.0001**[d] | **0.0002**[c] | 0.8742 | **<0.0001**[d] |
| KN4700 | **<0.0001**[d] | **0.0271a** | **<0.0001**[d] | **<0.0001**[d] | **<0.0001**[d] | 0.1307 |
| KN5088 | **<0.0001**[d] | 0.055 | **<0.0001**[d] | 0.2147 | 0.1724 | **0.0054**[b] |
| KN5092 | **<0.0001**[d] | **<0.0001**[d] | **<0.0001**[d] | **0.0026**[b] | **<0.0001**[d] | 0.2214 |
| KN5095 | **<0.0001**[d] | **0.0348**[a] | **<0.0001**[d] | 0.4445 | 0.077 | 0.5444 |
| KN1850 | **<0.0001**[d] | **0.0076**[b] | **<0.0001**[d] | **0.026**[a] | 0.7468 | **0.005**[b] |
| KN0543 | **<0.0001**[d] | **0.0211**[a] | **<0.0001**[d] | 0.5378 | 0.1996 | 0.7633 |
| KN3824 | **<0.0001**[d] | **<0.0001**[d] | **<0.0001**[d] | **<0.0001**[d] | 0.4834 | **<0.0001**[d] |

Statistically significant P values are marked in bold.

[a] – P value < 0.05, [b] - P value < 0.01, [c] – P value < 0.0005, [d] – P value < 0.0001
